# Bidirectional interactions between gut microbiota and fluorochemical biotransformation and bioactivity

**DOI:** 10.64898/2026.05.15.725488

**Authors:** Maja Stevanoska, Jorge Peña-Díaz, Mara L. Bieler, Raúl Fernández Cereijo, Jacob Folz, Leah Gächter, Silke I. Probst, Nika Sokolova, Serina L. Robinson, Nicholas A. Bokulich, Shana J. Sturla, Georg Aichinger

## Abstract

Fluorinated chemicals are increasingly prevalent in pharmaceuticals and agrochemicals, yet their influence on the human gut microbiome and the potential for microbial biotransformation to alter therapeutic and toxicological profiles remain poorly understood. Here, we investigated the bidirectional relationship between 15 structurally diverse fluorinated chemicals and the gut microbiota by using an *ex vivo* high-throughput fermentation system. Screening revealed that flutamide, fluazinam, and pretomanid were consistently biotransformed across the donor microbiomes, while other compounds showed substantial inter-individual variability in degradation. Furthermore, exposure to fluorinated chemicals induced compound-specific shifts in microbial diversity and community composition, demonstrating their capacity to alter gut microbial ecology. Using a computational workflow combining *in silico* biotransformation predictions with untargeted LC-MS/MS analysis, we identified nitroreduction as the primary gut microbial transformation across all three compounds. Single-strain experiments confirmed that the nitroreduction of flutamide to flu-6, previously attributed only to hepatic metabolism, is a widespread capacity among gut bacterial strains. Finally, *in vitr*o cytotoxicity assays and *in silico* modelling further revealed flu-6 to be a less hepatotoxic derivative than the parent compound, suggesting a potential detoxifying role for the gut microbiota. Together, these findings establish an integrated *ex vivo*, *in vitro*, and *in silico* approach for assessing the bidirectional interactions between fluorinated chemicals and the gut microbiome.

## 1. Introduction

The use of synthetic fluorinated pharmaceuticals and agrochemicals has steadily increased over the past decades, and currently more than 20% of pharmaceuticals^1^ and 50% of agrochemicals^2^ contain at least one fluorine atom. This expansion is driven by the favorable physicochemical properties associated with fluorination,^3,4^ together with the widespread incorporation of fluorine to enhance chemical and metabolic stability, which has contributed to the environmental and biological persistence of numerous fluorinated compounds. This persistence has reinforced the prevailing perception that fluorinated chemicals are broadly resistant to microbial metabolism and biodegradation. While the human microbiota harbors enzymes capable of cleaving carbon–fluorine bonds,^5^ other biotransformation steps, such as reduction, hydrolysis, conjugation, or nitroreduction, are more prevalent and favorable.^6^ Yet, the extent of these gut microbial interactions remains poorly understood, highlighting a need for systematic studies to refine our understanding of how gut microbiota transform diverse fluorinated pharmaceutical and agrochemical structures, as well as the reciprocal impact of these compounds on the gut microbiota.

Microbial transformations of fluorinated chemicals have the potential to alter therapeutic and toxicological profiles of these molecules. For example, *Escherichia Coli* increase the cytotoxicity of the fluorinated antineoplastic agent fludarabine through hydrolytic activation, yielding halogenated purine bases.^7^ Similarly, fluoropyrimidines such as 5-fluorouracil (5-FU) are metabolically inactivated by gut bacteria via reduction of the pyrimidine ring, a pathway shared with host metabolism.^8^ Several of the resulting metabolites, including fluorocitrate, α-fluoro-β-alanine, and fluoroacetate, disrupt the citric acid cycle and have been linked to neurotoxicity in cats^9^ and cardiotoxicity in humans.^10^ In contrast, although agrochemicals such as fluometuron and flubendiamide undergo microbial degradation in soil,^11,12^ there is currently no evidence of their degradation by human gut microbiota. Thus, gut microbial metabolism of fluorinated chemicals can generate biologically active metabolites, but their identity, extent, and inter-individual variability in metabolic capacity remain poorly studied. Beyond direct xenobiotic metabolism, exposure to fluorinated chemicals may also perturb the gut microbiome composition, which could in turn have consequences for the microbial metabolic potential and host-microbe interactions. Although microbiota shifts in response to specific fluorinated compounds have been reported previously,^8,13^ more information is needed to identify and predict how specific fluorinated chemical classes disrupt the gut microbiome, and consequences thereof for host health. A critical step to bridge this gap is the systematic evaluation of the bidirectional interactions linking microbial transformation of these compounds with the chemically induced changes in community structure and function across diverse fluorinated compounds under controlled, physiologically relevant conditions.

In this study, we evaluated 15 commonly used fluorinated pharmaceuticals and agrochemicals, examining their susceptibility to gut microbial transformation, associated inter-individual variability, and effects on microbial composition. We performed e*x vivo* fecal fermentation experiments using stool samples from healthy human donors under physiological, colon-like conditions. Chemical depletion was monitored by LC–MS/MS, integrating targeted quantification to track chemical degradation and untargeted analysis to profile microbially derived transformation products. Computational metabolism prediction tools were applied to support metabolite annotation, while QSAR modeling was used to explore the potential toxicological relevance of selected microbial metabolites. Together, these findings enabled us to identify microbiome-associated transformations, prioritize candidate metabolites for follow-up confirmation, and demonstrate a clear bidirectional relationship between fluorinated chemicals and the gut microbiome.

## 2. Materials and Methods

### 2.1. Chemicals and reagents

Capecitabine, pretomanid, fulvestrant, letermovir, posaconazole, ezetimibe, tembotrione, fluopicolide, elagolix, and fluazinam were purchased from Merck (Switzerland). Aprepitant, acetonitrile, mass spectrometry (MS)-grade methanol, and formic acid were purchased from Fisher Chemical. Flutamide and ciprofloxacin were purchased from Thermo Fisher Scientific (Switzerland).

Flu-6 was purchased from Sigma Aldrich. Lasmiditan and fluometuron were purchased from Chemie Brunschwig. Chemical stocks were diluted in DMSO and were stored at -20°C until analysis. The ingredients and vendor information for basal yeast casitone fatty acid (bYCFA) growth medium and its supplementation with a complex mixture of carbohydrates and mucin carbon sources (6C+Muc) are listed in the Supplementary Table S1.

### 2.2. Human stool collection

Stool samples were obtained from healthy volunteers after written informed consent was provided, confirming fulfillment of the inclusion criteria. The study was reviewed and approved by the ETH Zürich Ethics Commission under the proposal EK 2024-N-161. Eligible donors had regular bowel movements (between once every three days and up to three times per day), no history of chronic inflammatory bowel disease, no use of immunosuppressants, anticoagulants, or medications affecting gut transit or digestion during the month preceding donation, no recurrent intestinal discomfort, and no prior surgical interventions on the intestine. Samples were collected anonymously, with donors contributing a single sample into a sealable container (1 L) equipped with an AnaeroGen™ bag to establish anaerobic conditions within 30 seconds of collection. Samples were processed within three hours in an anaerobic chamber (5% H₂, 10% CO₂, 85% N₂). Fresh stool samples were immediately used for the *ex vivo* fermentation assay.

### 2.3. *Ex vivo* fermentation assay

The gut microbial biotransformation of 15 fluorinated chemicals was evaluated under simulated colon conditions using an *ex vivo* fermentation assay as described by Zünd et al.^14^ Experiments were conducted in an anaerobic chamber in an atmosphere of 10% CO_2_, 5% H_2_, and 85% N_2_ (Coy Laboratory Products Inc., Grass Lake, MI). To maintain strict anaerobic conditions, all reagents were placed in the chamber for pre-reduction for at least 24 hours. On the day of the experiment, the pH of the bYCFA medium was adjusted to 6.5 with 3 M HCl. Plates were prepared by aliquoting 1.5 mL of the pH-adjusted bYCFA medium supplemented with 0.25 mL of carbon sources (6C+Muc) into each well. All chemical treatments were diluted in water as required and added to the respective wells to a final concentration of 20 μM and a final volume of 2 mL per well. An equivalent amount of DMSO was used as the vehicle control. Finally, for each donor, fresh stool (1 g) was resuspended in 10 mL PBS, serially diluted to 10⁻□, and 20 µL of the respective slurry was inoculated into each well.

Immediately after inoculation, 40 µL aliquots from each technical replicate were pooled into a separate plate to serve as the initial timepoint (t_0_). From these pools, 20 μL were transferred to 96-well plates pre-filled with 180 μL ice-cold methanol for metabolomic analysis. Plates were then heat-sealed with aluminum microplate seals (Sigma Aldrich) and stored at −80 °C until further analysis. After 48 h of incubation, 100 μL from each well was transferred to a flat-bottom 96-well plate for optical density measurement (OD₆₀₀). Wells with divergent OD₆₀₀ values were excluded from pooling. Remaining technical replicates were pooled as described above. Finally, aliquots from these 48-hour pools were collected for metabolomics and DNA sequencing. Aliquots designated for DNA sequencing were first centrifuged for 20 minutes at 5500 rpm and 4°C, and the supernatant was discarded before storage. Plates and cell pellets were sealed and stored at −80 °C until further analysis.

Additionally, based on the initial screening results, a follow-up ex vivo experiment was performed on the most promising compounds concerning observed average biodegradation, using a higher density of feces (5% *w/w* in the incubation suspension) that was previously found optimal for quantifying microbial biotransformation under realistic physiological conditions.^15^ The protocol was the same as described above and was carried out in a Baker Ruskinn Concept 500 anaerobic chamber.

### 2.4. Liquid chromatography-tandem mass spectrometry (LC-MS/MS)

For LC-MS/MS analysis, samples were centrifuged at 2850 × g and 4 °C for 30 minutes. After centrifugation, 100 µL of supernatant from each well was transferred to 96-well LC-MS/MS plates (SureSTART WebSeal). The plates were then resealed and maintained at 4 °C in the autosampler. Every 20 injections, a blank, a quality control, and a calibration standard were included in the sequence. Samples were analyzed on a Vanquish HPLC system (Thermo Fisher Scientific) coupled to an ID-X Tribrid mass spectrometer (Thermo Scientific). Chromatographic separation was achieved with a Synergi Polar-RP column (4 µm, 80 Å, 30 mm × 2 mm) equipped with a SecurityGuard cartridge (Phenomenex) maintained at 40 °C. The mobile phases consisted of LC-MS-grade water containing 0.1% formic acid (A) and acetonitrile containing 0.1% formic acid (B), with a flow rate of 0.7 mL/min. The gradient elution program was set as follows: 5% B from 0–0.1 min, linear increase to 100% B from 0.1–1.8 min, hold at 100% B from 1.8–2.35 min, decrease to 5 % B from 2.35–2.4 min, and equilibration at 5% B from 2.4–2.8 min, followed by a 30 s needle wash before the next injection. Samples were analyzed in both positive (+3500 V) and negative (−2500 V) electrospray ionization (ESI) modes. Ion source settings were as follows: gas mode, static; sheath gas, 60 (arb. units); auxiliary gas, 15; sweep gas, 2; ion transfer tube temperature, 350 °C; vaporizer temperature, 400 °C. Full MS1 scans were acquired on the Orbitrap mass analyzer at a resolution of 60,000 over an *m/z* range of 60–900, with the RF lens set at 55%, and centroid data recorded. Targeted MS2 scans of the compounds of interest and internal standards were performed in the ion trap using optimized HCD or CID fragmentation settings (Supplementary Table S2). Seven iterative exclusion DDA runs were performed on pooled quality control samples at the end of acquisition. For DDA, two MS2 spectra were acquired per MS1 scan, with a dynamic exclusion duration of 2.5 s.

### 2.5. Untargeted metabolomics Data Processing

Untargeted LC–MS/MS data were processed using MS-DIAL (version 4.9.13), with detailed processing parameters provided in Supplementary Table S3. Metabolite annotation was based on accurate mass-to-charge (*m/z*) ratios and MS/MS spectral similarity to a combined reference library comprising the MassBank of North America (massbank.us) and NIST23 databases. Data acquired in positive and negative ESI modes were analyzed independently. Features were excluded if the maximum sample intensity was less than ten times the average signal observed in blanks. Peak height data were exported from MS-DIAL for further analysis.

### 2.6. Metabolite annotation and chemical transformation prediction

The canonical SMILES strings of the selected fluorinated chemicals were input into the metabolism prediction tools MicrobeRX^16^ and Pickaxe^17^ using one-step reaction predictions, excluding any subsequent transformations of the predicted metabolites. The predicted structure SMILES strings were converted to corresponding molecular formulas and monoisotopic masses. From these monoisotopic masses, *m/z* values for 45 potential ESI-MS adducts were calculated. MS-DIAL output files from positive- and negative-ionization mode analyses were consolidated for feature comparison and annotation. Measured *m/z* values of detected features were compared with calculated *m/z* values of predicted metabolites, and potential matches were retained for *m/z* differences of ≤ 0.005. To assess the biological plausibility of each match, features were grouped by experimental condition. Fold changes and Mann-Whitney U tests were calculated to compare treatment and control groups. Time-course profiles of the parent compound and each corresponding feature were plotted to visually evaluate time-dependent formation of the feature. Features showing parent-compound depletion accompanied by an increase in signal were considered metabolite candidates. To unequivocally confirm the identity of the flutamide metabolite flu-6, a synthetic flu-6 standard was purchased and analyzed using a targeted LC-MS/MS method. A complete list of matches, including SMILES codes and statistical results is provided in Supplementary Table S4, the reduction of candidate numbers with data processing steps in Supplementary Table S5.

### 2.7. Cell culture and cytotoxicity assays

The cytotoxicity of flu-6 was evaluated against flutamide using a cell viability assay. HepG2 cells (human hepatocellular carcinoma cell line) were maintained in Dulbecco’s Modified Eagle Medium (DMEM) supplemented with 10% (v/v) fetal bovine serum (FBS) and 1% (v/v) penicillin–streptomycin. Cells were cultured for two weeks prior to experiments in a humidified incubator at 37 °C with 5% CO₂. For viability assays, HepG2 cells were seeded in 96-well plates at a density of 14,000 cells per well in 100 µL of DMEM. Cells were grown to confluence for 96 h and then exposed to flutamide or flu-6 at concentrations ranging from 25 to 200 µM. Triton X-100 (0.1%) and DMSO (0.1%) were included as positive and vehicle controls, respectively. After 4 or 24 hours of exposure, the media was then removed, and the cells were washed with phosphate-buffered saline (PBS). Fresh medium (45 µL) and 50 µL of the CellTiter-Glo reagent were added to each well. Plates were agitated for 2 min on an orbital shaker and incubated in the dark for 10 min. Subsequently, 75 µL of the reaction mixture was transferred to white half-area 96-well plates, and luminescence was measured using a microplate reader (Infinite M200 Pro, Tecan Group, Männedorf, Switzerland).

### 2.8. Physiologically-based kinetic (PBK) modeling

A previously developed human, microbiome-competent PBK model^18^ was adapted to predict the impact of microbial biotransformation on flutamide pharmacokinetics and systemic concentrations of microbially formed flu-6. Minor modifications to the model concept included the shift from female to unisex physiology, the addition of a stomach compartment for a more accurate representation of gastrointestinal transit, and a simplification by including gut tissue into the “portal vein perfused tissue” compartment. Coefficients for blood/tissue partitioning and the fraction unbound in plasma were calculated using the algorithms of Berezhkovskiy and Lobell & Sivarajah, respectively, as provided by qivivetools,^19^ based on LogP values for flutamide (3.3) and flu-6 (2.8) as computationally predicted by the XLOGP3 tool.^20^ Parameters for flutamide’s gastrointestinal absorption and hepatic biotransformation were taken from van Tongeren et al.^21^ Caco-2 permeability of flu-6 was predicted using the algorithm of Kamiya et al.^22^ and scaled to *in vivo* absorption using the algorithm of Sun et al.^23^ Gut microbial flu-6 formation rates were estimated from fecal fermentation assays by comparison to a standard curve and scaled to *in vivo* biotransformation rates by fecal mass. The model was compiled in Berkeley Madonna and evaluated by predicting plasma flutamide levels within a 2-fold margin of in vivo pharmacokinetic data.^24^ The model code, along with all parameters, is available via Zenodo^25^ (https://doi.org/10.5281/zenodo.19682173), predictions used for model evaluation are included in the Supplementary Figure S1.

### 2.9. Single-strain confirmation of flutamide reduction

To confirm the biotransformation of flutamide, the following bacterial strains were obtained from DSMZ: *Lacticaseibacillus rhamnosus* DSM111733, *Enterococcus faecium* DSM17050, *Schaalia odontolytica* DSM1912, *Gordonibacter pamelaeae* DSM110924, *Escherichia coli* K12 DSM18039 and *Streptococcus salivarius* subsp. salivarius DSM20067. These strains were cultured anaerobically overnight in Gut Microbiota Medium (GMM) and then diluted to an OD_600_ of 0.05 in triplicate in 2 mL polypropylene tubes to the final volume of 1 mL. Fluorinated chemicals were added individually at a final concentration of 200 µM. 250 µL aliquots were sampled from each culture at 0, 24, and 48 h post-inoculation. Samples were immediately added to 750 µL of ice-cold acetonitrile/methanol (1:1 v/v) and centrifuged for 10 min at 4000 rpm. The supernatants were transferred to glass vials and analyzed by HPLC and LC-MS/MS.

Flutamide biotransformation was quantified using HPLC-DAD (UV-vis) (Ultimate HPLC system, Dionex). HPLC experiments were conducted using solvents A (water + 0.1% formic acid) and B (acetonitrile + 0.1% formic acid). Chromatographic separation was achieved with a Nucleoshell RP 18plus column (particle size 5 μm, 150 × 3 mm, Macherey-Nagel) using a 10 µL injection volume. The flow rate was maintained at 1 mL/min with a linear gradient as follows: 10% B for 2 min,10-90% B over 8 min, 90-10% B over 1 min, 10% B for 0.5 min. Absorbance peak areas were measured at 305 nm and integrated using Chromeleon 7.2.10. Integrated peak areas were converted to concentrations in µM using calibration plots built with authentic standards of flutamide and flu-6.

The samples were further analyzed with liquid chromatography coupled to high-resolution tandem mass spectrometry (LC-MS/MS). The LC-MS/MS method was adapted from Yu *et al.*^26^ The samples were separated over a Waters ACQUITY Premier BEH C18 column (1.7 μm, 100 × 2.1 mm) at 30 °C with a concentration gradient (solvent A: water + 0.1□% formic acid, and solvent B: methanol + 0.1□% formic acid) at a flow rate of 0.2 mL/min (10 μL injections). The following gradient was used: 5□% B for 1.5 min, 5-95□% B over 6 min; 95□% B for 2 min; 5□% B for 2 min. Spectra were acquired in full-scan mode over *m/z* 100 − 1000 at a resolution of 70,000. The acquired data were analyzed using MZmine 4.0.3.^27^

### 2.10. DNA purification, amplicon library preparation and sequencing

Cell pellets from 48 h *ex vivo* fermentations were resuspended in 500 µL of ultrapure water, and a 200 µL aliquot was used for DNA extraction with the MagMAX Microbiome Ultra Nucleic Acid Isolation Kit on the KingFisher Apex platform (Thermo Fisher Scientific), according to the manufacturer’s instructions. The ZymoBIOMICS Microbial Community Standard was used as positive control, and molecular-grade water as negative control on each plate. Bead beating was performed for 5 min at 30 Hz. DNA was eluted in 50 µL of elution buffer. DNA concentrations were measured using the Qubit dsDNA Quantification Assay Kit (Thermo Fisher Scientific), following the manufacturer’s instructions. DNA was stored at −20 °C until further processing.

Amplicon libraries for the 16S rRNA gene were prepared using the HighALPS ultra-high-throughput protocol.^28^ The 16S rRNA gene V4 region was amplified using primers 515F (5′-GTGYCAGCMGCCGCGGTAA-3′) and 806R (5′-GGACTACNVGGGTWTCTAAT-3′).^29,30^ PCR reactions contained 1.5 µL of template DNA and were amplified using KAPA HiFi HotStart ReadyMix (Roche) according to the manufacturer’s instructions. Cycling conditions were as follows: initial denaturation at 95 °C for 5 min; 28 cycles of 98 °C for 20 s, 55 °C for 20 s, and 72 °C for 25 s; and a final extension at 72 °C for 5 min. Amplicons were pooled at equimolar concentrations and purified using Agencourt AMPure XP magnetic beads at a 0.7× bead-to-sample ratio (Beckman Coulter). Library quality was assessed using a TapeStation system (Agilent Technologies). The final pooled library was sequenced at the Functional Genomics Center Zürich (FGCZ) using 300-bp paired-end reads on an Illumina NextSeq 2000, with a 20% PhiX spike-in.

### 2.11. Microbial diversity and statistical analysis

Sequencing data were analyzed in QIIME 2.^31^ Paired-end reads were processed with DADA2,^32^ and reads were truncated at 240 bp in both directions to remove low-quality regions, primers, and adapters, and a minimum fold-parent-over-abundance threshold of 4.0 was applied. Taxonomic classification was performed using a pre-trained, human–stool–weighted Naïve Bayes classifier^33–35^ trained on SILVA 138.^36^ Reads assigned to mitochondria, chloroplasts, archaea, and eukaryotes were removed.

For diversity analysis, data were rarefied to 1000 reads per sample. Alpha diversity (Shannon) was analyzed using Linear Mixed Models (LMM) with treatment as a fixed effect and donor as a random intercept. Differences relative to vehicle control were assessed using Dunnett’s correction for multiple comparisons. Beta diversity was evaluated by calculating Bray-Curtis dissimilarity between treatment samples and their paired vehicle controls. Significance was assessed using a pairwise PERMANOVA stratified by donor with P-values adjusted for multiple comparisons. Differential abundance analysis was performed at the genus level using ANCOM-BC2.^37^ To account for the repeated measures, treatment was modelled as a fixed effect and donor as a random intercept. Differences in genus abundance between each treatment and the vehicle control were assessed using Dunnett’s test, with P-values adjusted for multiple comparisons using Holm’s method.

## 3. Results

### 3.1. Microbial biotransformation and community-modulating effects of fluorinated chemicals on the human gut microbiome

To investigate bidirectional interactions between gut microbiomes and fluorinated chemicals, we performed high-throughput *ex vivo* fecal fermentations using stool from 13 healthy donors. This approach allowed us to simultaneously quantify the depletion of fluorinated chemicals by targeted LC-MS/MS and characterize their effects on microbial community dynamics by 16S rRNA sequencing after 48 h of incubation. Incubations without fecal slurry (abiotic controls) were included to differentiate microbial from non-microbial degradation. Results revealed that concentrations of flutamide, fluazinam, and pretomanid were consistently reduced across all donor microbiome samples (Figure 1). In contrast, other compounds, such as capecitabine and fulvestrant, showed high interindividual variability, with depletion observed only in a subset of donor communities. Unexpectedly, posaconazole did not follow these patterns and instead showed an apparent increase in concentration after 48 h relative to the baseline; this likely represents an artifact driven by matrix effects or by increased drug solubility over the course of the fermentation.

**Figure 1.**
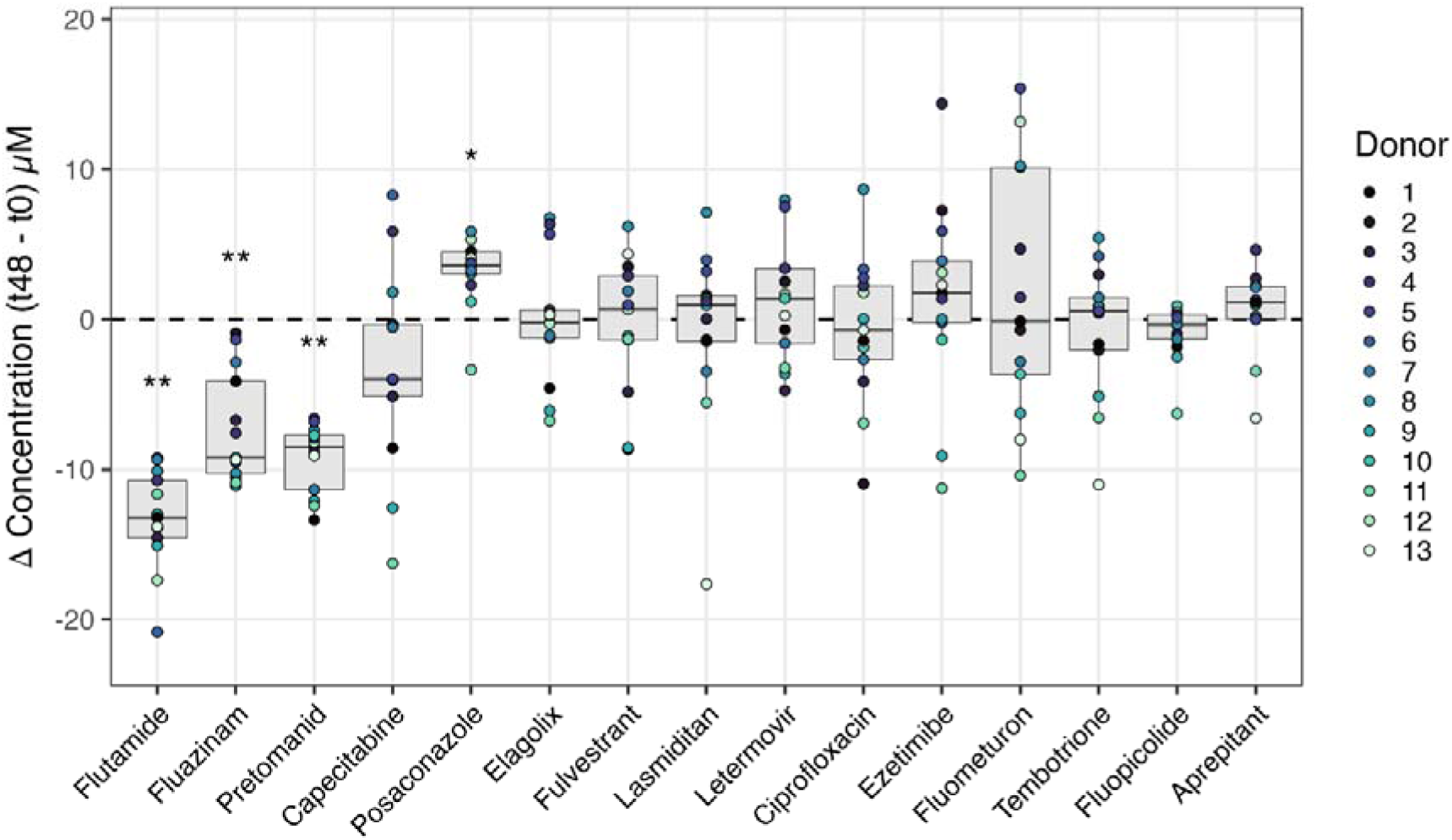
Depletion of fluorinated compounds in *ex vivo* stool cultures. Evaluation of 15 fluorinated chemicals screened in an *ex vivo* fermentation assay derived from human stool samples. Boxplots represent the change in concentration between inoculation (t0) and 48 h across individual donors (n = 13). Statistical significance was assessed using a one-sample Wilcoxon signed-rank test with Holm’s correction for multiple testing. Statistical significance is reported as follows: * *p* ≤ 0.05; ** *p* < 0.01.

Having quantified chemical depletion, we next evaluated how these fluorinated compounds affected overall microbial community composition. Alpha diversity analysis revealed distinct treatment responses, characterized by significant reductions in the Shannon index following exposure to fluazinam, pretomanid, fluopicolide and ciprofloxacin (Figure 2A). Despite significant compound depletion, flutamide exposure resulted in only modest changes in alpha diversity compared with the vehicle control. Evaluation of beta diversity further supported these observations, with fluazinam, pretomanid, and ciprofloxacin inducing the most pronounced shifts in overall community composition with a higher Bray-Curtis dissimilarity relative to the vehicle control (Figure 2B).

**Figure 2.**
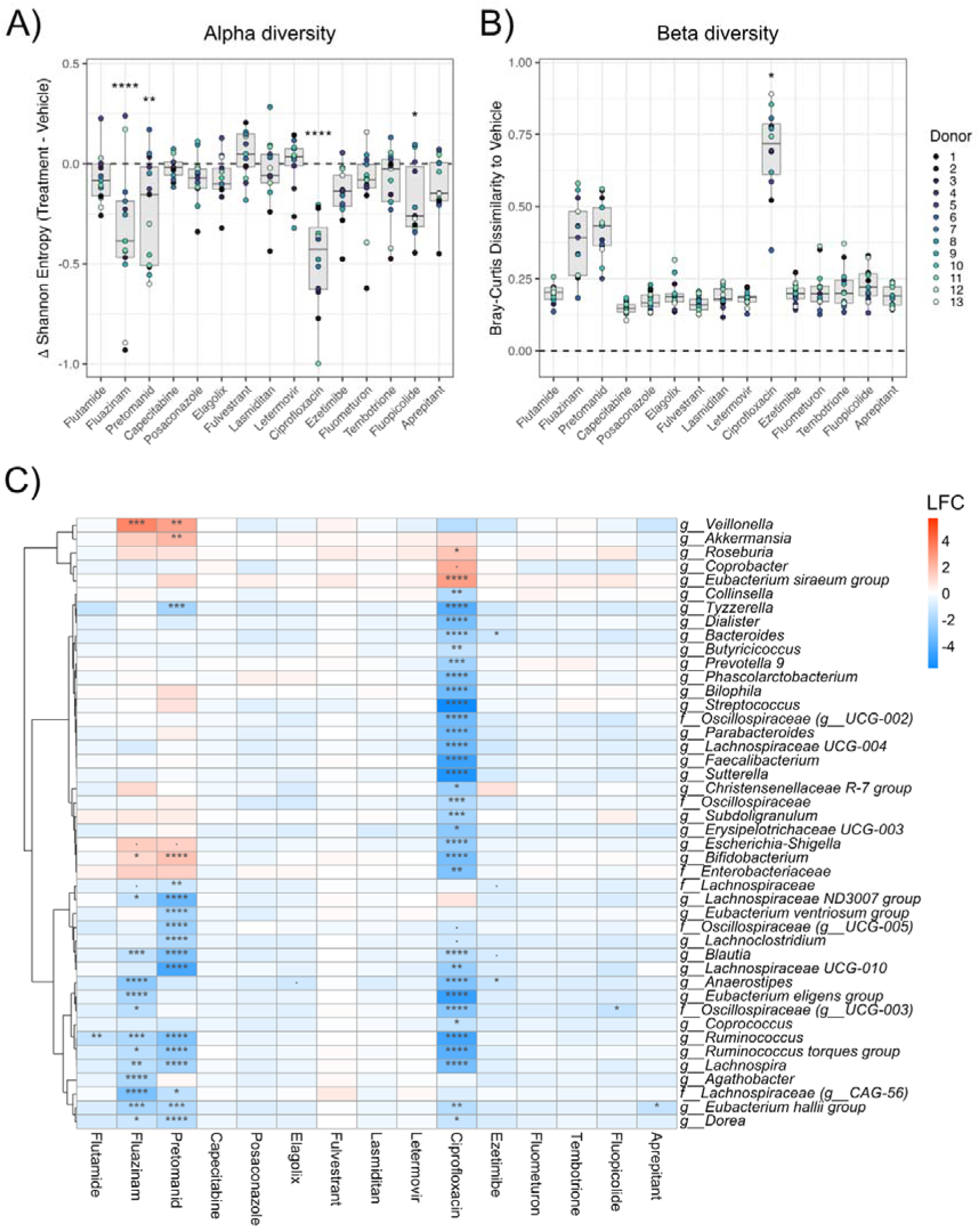
Fluorinated compounds differentially alter gut microbial diversity and community composition. **A)** Changes in alpha diversity (Shannon), calculated as the difference between each treatment and the vehicle control. Significant differences were determined using Linear Mixed Models with Dunnett’s adjustment for multiple comparisons. **B)** Beta diversity (Bray-Curtis dissimilarity) between treatment samples and the corresponding paired vehicle control. Significance was assessed using pairwise PERMANOVA stratified by donor, followed by Benjamini-Hochberg (FDR) adjustment for multiple comparisons. Dashed lines (y=0) indicate the vehicle baseline. **C)** Differential abundance of bacterial genera at 48 h post-inoculation assessed using ANCOM-BC2. Only taxa with a significant differential abundance (p < 0.05) in at least one treatment are shown. Dunnett’s test was used to compare each treatment against the vehicle control, with Holm’s adjustment for multiple comparisons. Colors represent the log-fold change (LFC), where red indicates enrichment and blue indicates depletion relative to the vehicle. Statistical significance, based on adjusted *p*-values, is reported as follows: *p* < 0.10; * *p* ≤ 0.05; ** *p* < 0.01; *** *p* < 0.001; **** *p* < 0.0001.

To identify which specific community members were depleted or enriched following exposure to the fluorinated chemicals, we performed differential abundance analysis using ANCOM-BC2 (Figure 2C). As expected, ciprofloxacin reduced the relative abundance of a wide range of bacterial genera. Fluazinam and pretomanid also altered bacterial community structure by significantly reducing the abundance of multiple microbial taxa, including the families *Lachnospiraceae, Eubacteriaceae,* and *Ruminococcaceae*. In contrast, these treatments were also associated with significant enrichment of the commensal genera including *Veillonella* and *Bifidobacterium*. This expansion may reflect the depletion of susceptible bacteria, thereby creating an ecological niche for less susceptible taxa to expand. Conversely, treatments such as flutamide were associated with fewer taxa that were differentially abundant. Thus, the significant depletion of flutamide (Figure 1) occurred without widespread disruption of the microbial community. These results point toward bidirectional interactions: while fluorinated compounds can exert selective pressure on specific bacterial taxa and alter community composition, the observed depletion of these compounds suggests a potential capacity for microbial biotransformation.

### 3.2. Identification of specific bacterial taxa associated with compound biotransformation

Having established that some fluorinated chemicals are depleted in a gut-mimicking environment, we next sought to identify the specific bacterial taxa associated with these biotransformations. To link microbial composition to the different levels of microbiome-driven depletion, compound concentrations were normalized relative to the abiotic control (Supplementary Figure S2) followed by a Spearman rank correlation between the centered log-ratio (CLR) transformed abundances of taxa and the normalized percentage of drug remaining after 48 h. Results revealed a wide range of microbe-compound associations. Positive correlations likely reflect taxa with reduced susceptibility, which allows them to persist or bloom after treatment. In contrast, negative correlations associate a higher abundance of specific taxa with the degradation of fluorinated compounds, suggesting either a potential role in their biotransformation (Figure 3A) or a fitness advantage following their removal. By focusing on the compounds with the highest degradation rates in Figure 1, we identified several bacterial candidates associated with compound removal. For instance, pretomanid depletion was strongly negatively associated with *Lachnospiraceae* (R = -0.76, *p*

**Figure 3.**
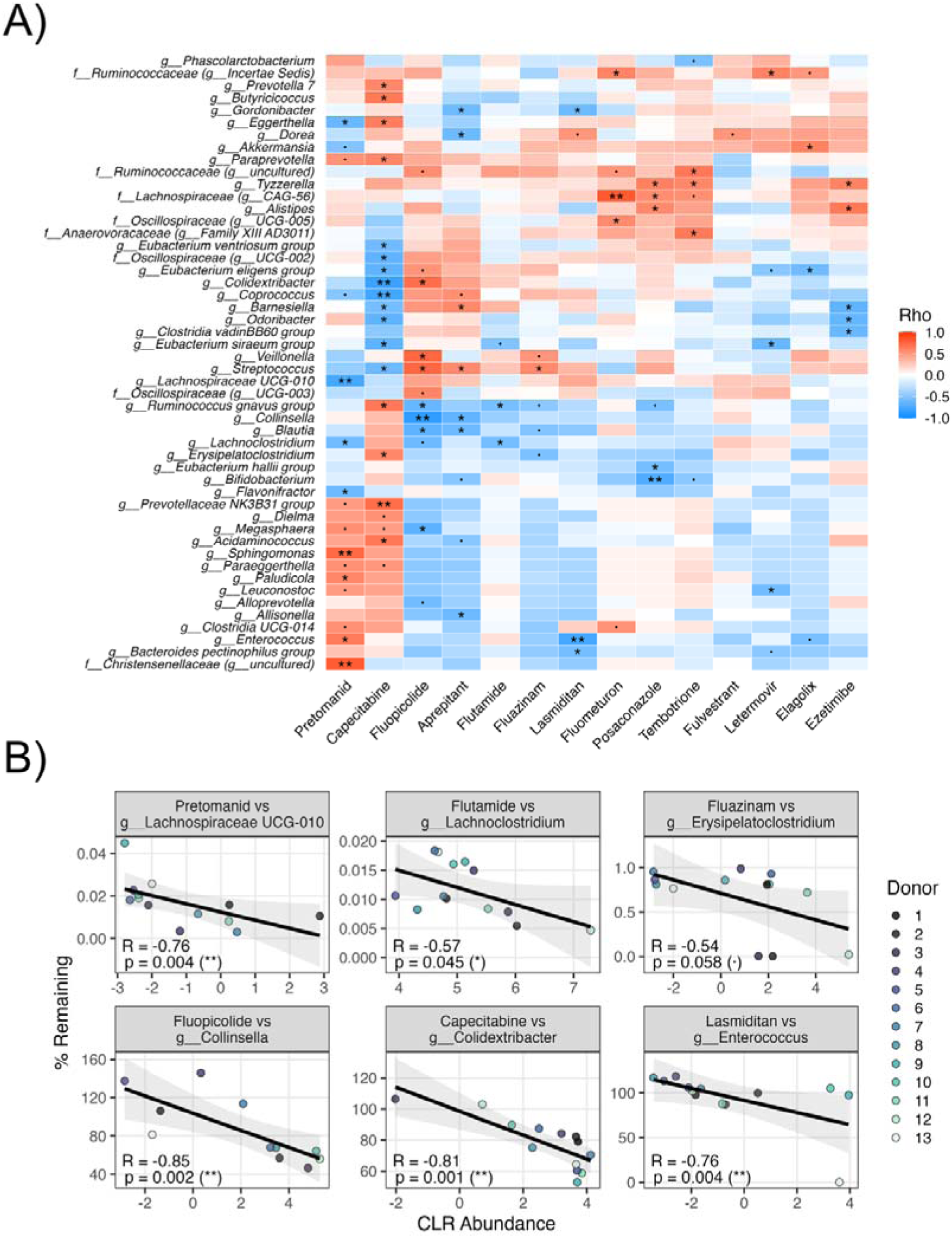
Associations between gut bacterial genera and drug depletion. **A)** Heatmap of Spearman rank correlation coefficients between centered log-ratio (CLR) transformed bacterial abundances and the normalized percentage of drug remaining at 48 h (Supplementary Figure S2). The heatmap shows the top 50 genera with the strongest correlation coefficient (R). Red denotes positive correlations, whereas blue denotes negative correlations. Ciprofloxacin was excluded from the correlation analysis. **B)** Scatter plots for representative treatments illustrating associations between bacterial abundance and fluorinated compound depletion. Points represent individual samples colored by donor. Black lines indicate linear regression with 95% confidence intervals. Statistical significance is reported as follows: · *p* < 0.10; * *p* ≤ 0.05; ** *p* < 0.01.

= 0.004), while flutamide and fluazinam showed moderate association with *Lachnoclostridium* (R = -0.57, *p* = 0.045) and *Erysipelatoclostridium* (R = -0.54, *p* = 0.058), respectively. Notably, the strongest correlations in the dataset were observed for compounds with high inter-individual variability, linking, for example, fluopicolide with *Collinsella* (R = - 0.85, *p* = 0.002), capecitabine with *Colidextribacter* (R = -0.81, *p* = 0.001), and lasmiditan with *Enterococcus* (R = -0.76, *p* = 0.004; Figure 3B). Positive correlations were also observed, for example, between fluazinam and *Streptococcus* (R = 0.57, *p* = 0.047), which could be driven by shifts in the abundance of other taxa or by reduced growth for resource allocation. These associations can also be confounded by antimicrobial activity. For example, treatments such as ciprofloxacin deplete a wide range of susceptible taxa, resulting in lower bacterial abundances when drug exposure remains high, leading to negative correlations (Supplementary Figure S3). Yet, maximum optical density (OD_600_) remained stable across treatments, suggesting no major change in total bacterial growth despite treatment-associated shifts in community composition (Supplementary Figure S4). While these statistical correlations cannot definitively establish causation, particularly since biotransformation is unlikely to directly drive bacterial growth, they provide a valuable strategy for hypothesis generation and for prioritizing candidate taxa for future mechanistic characterization.

### 3.3. Fluorinated chemicals undergo variable microbial transformation under simulated colonic conditions

To characterize the kinetics and donor-dependent variability of gut microbial biotransformation, we selected four chemicals that were most robustly depleted in our initial screen (Figure 1) and exposed them to fecal communities derived from a separate group of 10 individuals. To capture temporal dynamics, the cultures were sampled at 0, 6, 24, and 48 h post-inoculation. At each time point, we performed targeted and untargeted LC-MS/MS analyses to monitor compound removal and identify biotransformation products. For all donor samples, flutamide depletion began at 6 h and was nearly complete by 24 h, while concentrations in the abiotic control remained largely unchanged. Fluazinam was rapidly depleted across all microbiomes (within 6 h), whereas its decline was slower in abiotic controls. Pretomanid was removed more gradually in most donor samples, and was completely lost by 48 h, whereas it was minimally changed in abiotic controls. In contrast, capecitabine depletion was minor and comparable to the abiotic control, suggesting it was not microbially driven. Corresponding data are shown in Supplementary Figure S5.

To evaluate how colon-like microbial densities impact the biotransformation potential of fluorinated compounds, *ex vivo* fermentation assays were performed using a higher density of fecal microbes. To capture the rapid depletion kinetics, the cultures were sampled at finer resolution than the prior analysis (0, 1, 3, 6, 9, and 24 h). Flutamide and fluazinam were rapidly depleted (within 1 h) in most microbiomes (Figure 4A-B). In comparison, pretomanid was removed slower and was nearly completely depleted within 3 h in most donor microbiomes; however, 3 of the 10 donors exhibited reduced metabolic capacity, requiring 6-9 h for full removal (Figure 4C). Consistent with our previous results, capecitabine was largely resistant to biotransformation (Figure 4D). These findings demonstrate that physiologically relevant microbial densities dramatically accelerate the biotransformation of susceptible fluorinated drugs, while still maintaining distinct, donor-specific metabolic profiles.

**Figure 4.**
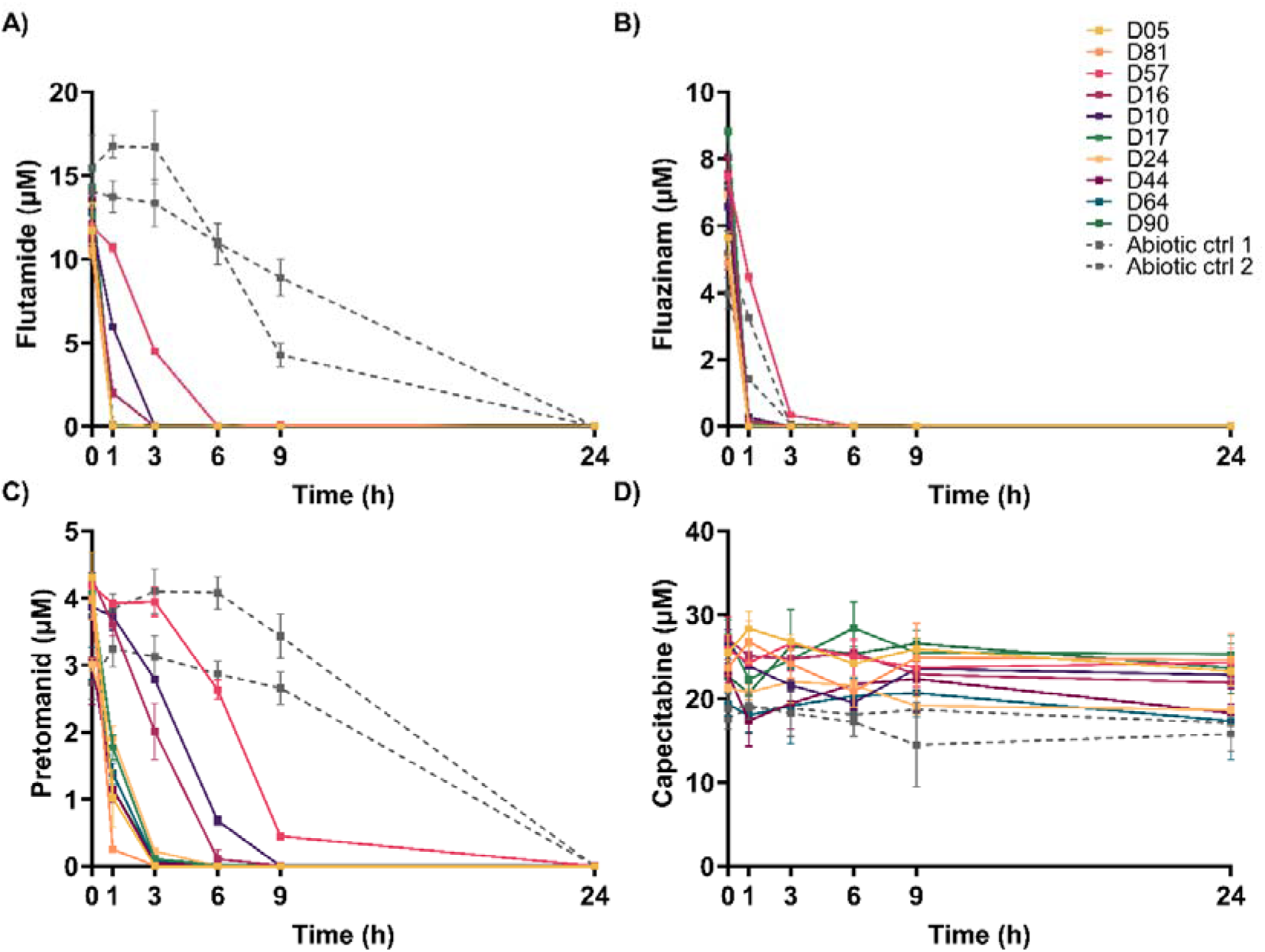
Kinetics of microbial metabolism of A) flutamide, B) fluazinam, C) pretomanid, and D) capecitabine over time. Fecal slurries from 10 donors were incubated with the same compound and sampled at 0, 1, 3, 6, 9, and 24 h to monitor degradation kinetics. Dashed lines represent the abiotic controls. Data represent the mean ± SD of three technical replicates.

### 3.4. Identification of microbial biotransformation products

Given that flutamide, fluazinam, and pretomanid were the only compounds consistently depleted in our previous assays, we focused on identifying their previously uncharacterized microbial biotransformation products. First, putative metabolic products were predicted *in silico*, yielding 471, 268, and 431 candidate structures for flutamide, fluazinam and pretomanid, respectively. These structures were subsequently queried against untargeted LC–MS data from fecal incubations using a computational identification pipeline, narrowing the results to 2, 3, and 5 candidates, respectively (Supplementary Table 5). These structures corresponded to nitro-reduced metabolites of the three compounds, as well as one hydrolysis product of fluazinam and ether hydrolysis products of pretomanid (Figure 5).

**Figure 5.**
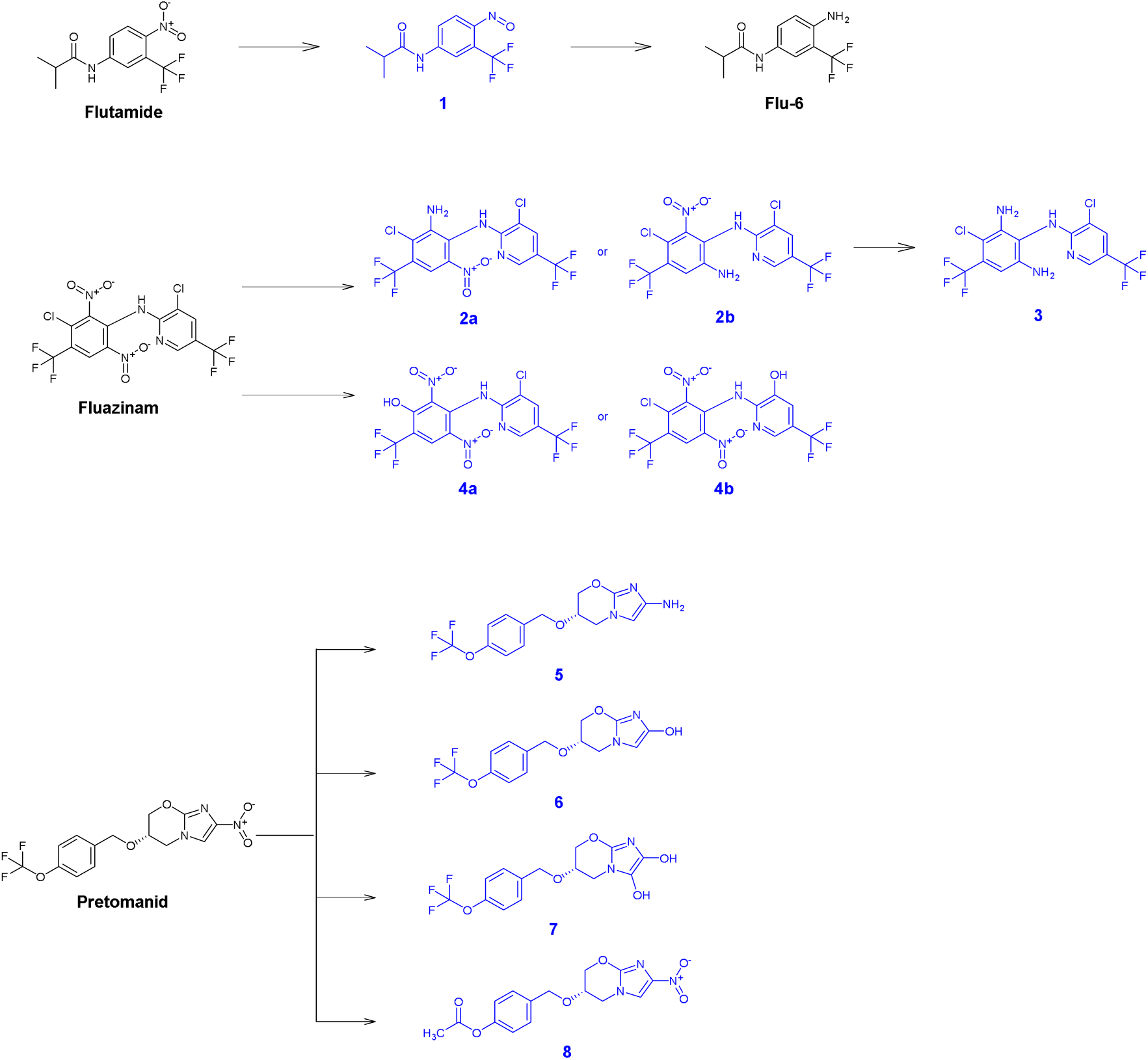
Chemical structures and proposed microbial degradation products for flutamide, fluazinam, and pretomanid. Chemical structures were predicted *in silico* using MicrobeRX and Pickaxe and matched with untargeted LC-MS data derived from *ex vivo* fecal fermentations. The identification confidence corresponds to putative structures (blue) and authentic chemicals standard confirmed metabolites (black).

Nitroreduction is a key transformation known to occur in the human gut microbiome, predicted here across all three compounds. In the case of flutamide, we observed the formation of a feature with *m/z* 247.10503, which was proposed as the amino product flu-6 (Figure 5A). In addition, a chemical with a mass consistent with the nitroso intermediate of the nitro reduction (*m/z* 259.06976, [M-H]^-^, compound **1**) was detected in low abundance and was clearly depleted over time (Supplementary Figure S6a). Since flu-6 was commercially available, this enabled the development of a targeted LC-MS/MS method to quantify flutamide degradation and flu-6 formation over 24 hours using the donor exhibiting the highest flu-6 biotransformation activity. Flu-6 levels reached approximately 1 µM (∼5% conversion) after 1 h, and remained relatively stable over 24 h (Figure 6). With this analysis, we confirmed the microbially driven nitro reduction of flutamide to flu-6, a compound known to form in the liver but not previously reported as a product of the human gut microbiota.

**Figure 6.**
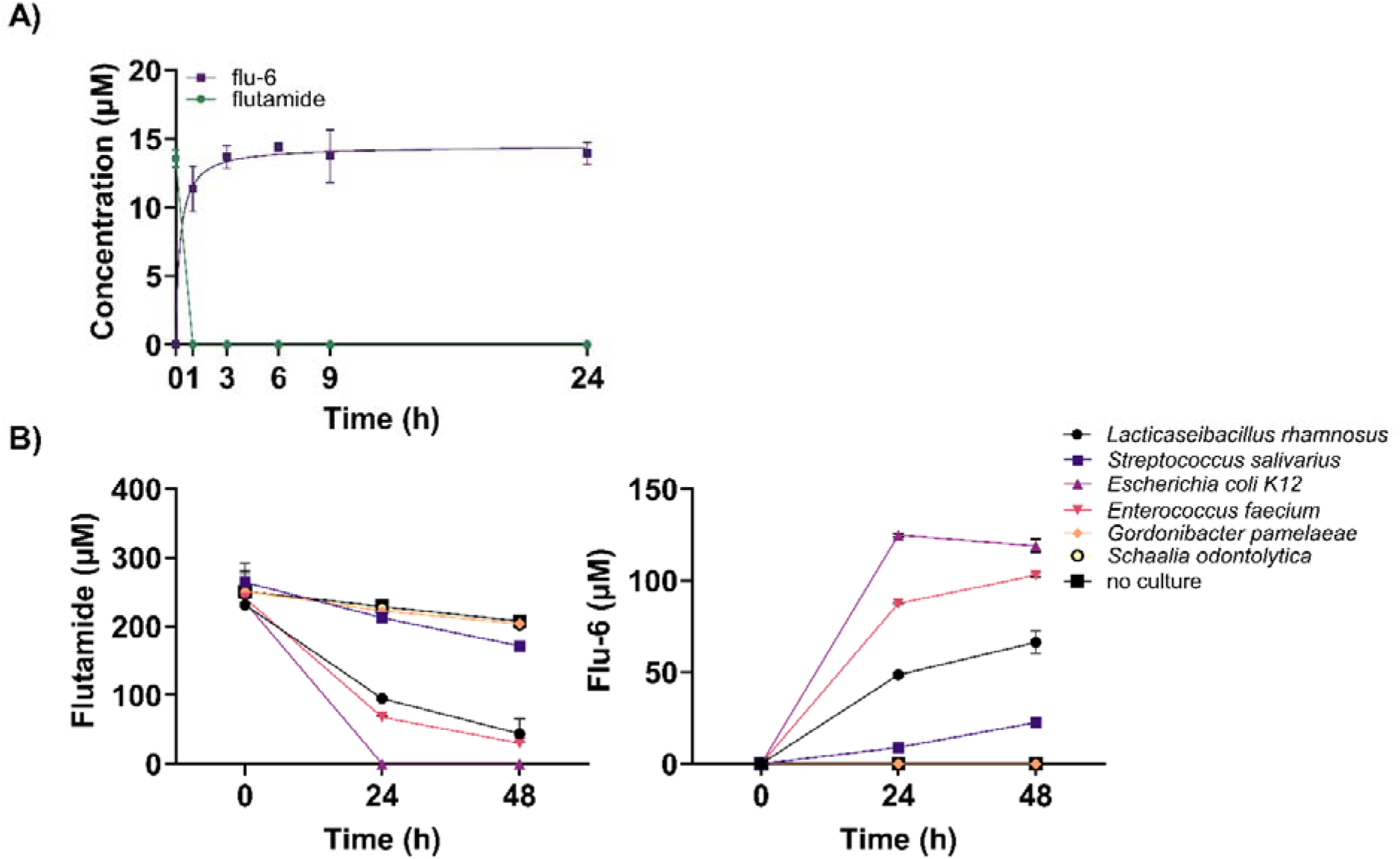
Biotransformation of flutamide to flu-6. A) Time-course biotransformation of flutamide to flu-6 in the donor with the highest biotransformation activity (donor 44). These data represent the mean ± SD of three replicates. B) Biotransformation of flutamide by *Streptococcus salivarius*. Depletion of flutamide (200 µM) and formation of flu-6 were quantified by HPLC at 0, 24 and 48 hours post-inoculation. Gut Microbiota Medium (GMM) was inoculated with *Streptococcus salivarius, E. coli*, *Enteroccocus faecium, Lacticaseibacillus rhamnosus, Schaalia odontolytica, Gordonibacter pamelaeae*, or left uninoculated as the abiotic control, and incubated anaerobically. The data represent the mean ± SD of three replicates.

**Figure 7.**
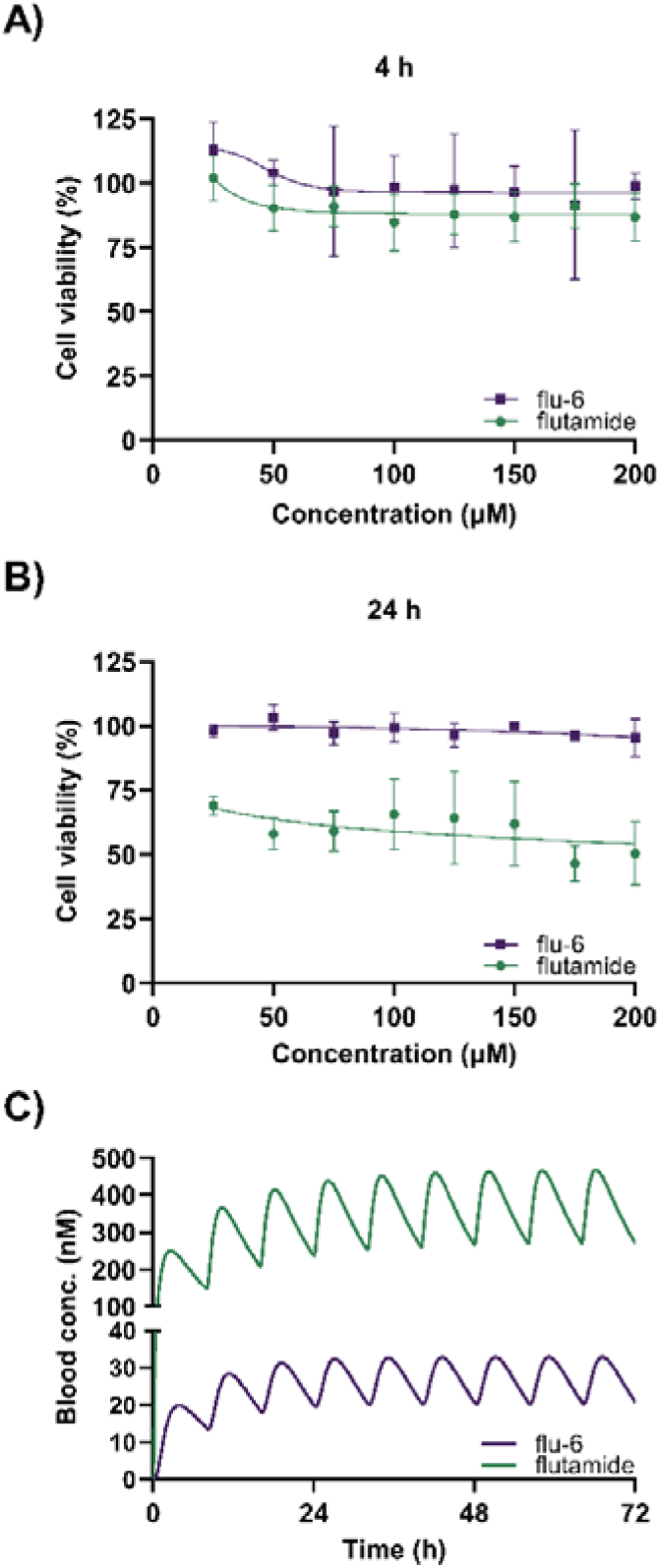
Cytotoxicity and estimated pharmacokinetics of flutamide and flu-6. Cell viability of HepG2 cells following exposure to increasing concentrations (25–200 µM) of flutamide or flu-6. Cells were treated for A) 4 or B) 24 h, and viability was assessed using the CellTiter-Glo^®^ assay. Data represent the mean ± SD of four biological replicates. C) Estimated blood concentrations of flutamide and microbially derived flu-6. Pharmacokinetics were predicted using a microbiome-competent PBK model under standard dosing (3x 500 mg/day).

Similar to flutamide, fluazinam also underwent nitroreduction, where we detected a mass consistent with the reduction of one of its two nitro groups (m/z 434.98462, [M+H]⁺, compound **2**, Figure 5B). Compound **2** accumulated initially but then declined after 3 h (Supplementary Figure S6b), suggesting that it was further transformed. Furthermore, we observed an additional ion with a mass consistent with the reduction of both nitro groups (*m/z* 405.01068, [M+H]^+^, compound **3**), which accumulated over time without further degradation. One last mass (*m/z* 444.97784, [M-H]^-^, compound **4**) was assigned to a hydrolytic dechlorination product, and the isotopic fingerprint supported the existence of only one chlorine atom in the structure, compared with two chlorines in fluazinam. Our method did not consider any fragmentation data for unknown chemicals; thus, discrimination between both pairs of isomers (Figure 5 2a-b and 4a-b) remains unresolved.

For pretomanid (Figure 5), three masses were consistent with predicted nitroreduction products. The first structure was predicted as the amino product of pretomanid nitroreduction (*m/z* 347.13138, [M+NH_4_]^+^, compound **5**). Compound **6** *(m/z* 348.13318, [M+NH_4_]^+^) was consistent with concomitant hydroxylation, and compound **7** (*m/z* 364.11185, [M+NH_4_]^+^) with a similar structure lacking the vicinal hydroxyl group. One additional mass was found matching compound **8** (*m/z* 332.08594, [M-H]^-^), likely a secondary hydrolysis degradation product.

### 3.5. Single-strain evidence of flutamide nitroreduction

Having established flu-6 as a microbially derived metabolite, we next sought to identify specific bacterial strains capable of performing this nitroreduction. For this, we screened six gut bacterial strains selected on the basis of taxonomic diversity, availability in the DSMZ collection, and prior evidence of growth in the presence of fluorinated compounds ^38,39^. Four of the six strains tested (*Streptococcus salivarius, E. coli*, *Enteroccocus faecium*, and *Lacticaseibacillus rhamnosus*) demonstrated the capacity to transform flutamide to flu-6. By the end of the 48-hour incubation, flutamide was completely depleted in the *E. coli* culture (35% *Streptococcus salivarius,* 87% *Enteroccocus faecium*, 81% *Lacticaseibacillus rhamnosus*), accompanied by a corresponding increase in flu-6 concentration (Figure 68). In contrast, flutamide remained mostly stable in both abiotic controls and cultures of the remaining two strains tested (*Schaalia odontolytica, Gordonibacter pamelaeae*), in which no flu-6 was detected (data not shown). These results suggest that nitroreduction of flutamide is a widespread capacity among several members of the gut microbiota.

### 3.6. An *in silico* and *in vitro* hazard characterization of flu-6

Given the high relevance of microbial nitroreduction and the confirmation of flu-6 formation from flutamide, the potential host impact of this transformation is of interest, particularly given the limited biological characterization of flu-6. To address this, we first used the Danish QSAR Database^40^ to predict potential human toxicity endpoints for flutamide and flu-6 (Supplementary Table S6). While both compounds were predicted to interact with androgen receptors (AR), inhibiting aromatase activity, and inducing cytochrome P450 3A4 (CYP3A4), a key enzyme involved in the metabolism of a large portion of drugs, flutamide had consistently higher probabilities for AR binding, antagonism, and inhibition (0.99, 0.95, and 0.94, respectively) compared to flu-6 (0.86, 0.52, and 0.77). Similar trends were observed for CYP3A4 induction (0.80 for flutamide vs. 0.65 for flu-6), deiodinase inhibition, and modulation of the sodium-iodide symporter, indicating broader and more potent bioactivity for flutamide. Both compounds scored similarly for liver cancer risk in rodent models (0.86 and 0.84), suggesting potential hepatotoxicity. Based on these predictions, biotransformation to flu-6 does not appear to substantially alter the toxicological profile of flutamide. However, given that hepatotoxicity has been reported as a clinical side effect of flutamide, hepatotoxicity was prioritized for experimental validation.

Given the potential for flu-6 hepatotoxicity, we evaluated its cytotoxicity in a human hepatic carcinoma cell line (HepG2) and compared it with flutamide. HepG2 cells were exposed to flutamide or flu-6 at concentrations ranging from 25 to 200 µM for 4 or 24 h, followed by assessment of cell viability. After a 4-hour exposure, neither flutamide nor flu-6 significantly altered cell viability at any of the tested concentrations (Figure 9A). However, after 24 h of exposure to 200 µM flutamide, cell viability decreased to approximately 50%. Yet, flu-6 did not induce substantial cytotoxicity under the same conditions, with no marked reduction in cell viability observed after 24 h exposure (Figure 9B). These data indicate that flu-6 is less cytotoxic to hepatic cells than flutamide, suggesting a potential detoxification role for the gut microbiota.

To translate these *in vitro* cytotoxicity results to the human context, we estimated the blood concentrations of flutamide and flu-6 using a microbiome-competent PBK model. In a standard 3x500 mg/day dosing scenario, steady-state maximal concentrations (C_max_) for both compounds were predicted to be achieved within three days, with flutamide reaching 450 nM and flu-6 reaching 33 nM (Figure 9C). These predicted concentrations are below the point of departure doses for in vitro cytotoxicity, supporting the limited hepatotoxic potential for flu-6 at physiologically relevant levels.

## 4. Discussion

This study demonstrates that exposure to certain fluorinated chemicals can elicit bidirectional interactions with the gut microbiome, with potential consequences for human health. Chemical exposures can drive specific compositional shifts within microbial communities, while the microbiota possesses the capacity to actively biotransform these compounds. Together, these findings highlight the critical role of the gut microbiota in modifying the chemistry, efficacy, and potential toxicity of fluorinated chemicals.

Gaining a better mechanistic understanding of the interactions between fluorinated chemicals and the human gut microbiota is crucial, given the increasing prevalence of these compounds, which now constitute nearly 50% of agrochemicals and 20% of pharmaceuticals approved in recent years.^1^ While it is well established that these compounds can drive taxonomic and functional changes,^38^ our results highlight the bidirectional nature of this interaction and reveal substantial inter-individual variability in biotransformation across donors, differences that carry potential downstream toxicological repercussions for the host. At the mechanistic level, the microbiota can interact with these compounds through both bioaccumulation and biotransformation. For instance, bacteria are known to bioaccumulate fluorinated pharmaceuticals and long-chain perfluorinated compounds, reaching intracellular concentrations of up to millimolar levels while maintaining growth.^41,42^ Biotransformation, in turn, relies on specific enzymatic activities; for example, taxa such as *Clostridia* and *Bacilli* encode enzymes capable of cleaving carbon-fluorine bonds in simple mono- and difluorinated compounds.^5^ However, for fluorinated compounds bearing -CF3 groups, C-F cleavage is generally unfavorable; as such, gut microbial enzymes preferentially catalyze other biotransformations, such as nitroreduction or conjugation, rather than defluorination.^6, 43^

To better understand these interactions, we screened 15 structurally diverse fluorinated chemicals and identified flutamide, fluazinam, and pretomanid as consistently biotransformed across donor-derived microbiomes (Figure 1). Upon exposure to fluorinated chemicals, we observed changes in microbial diversity and taxonomic structure, yet these community shifts were not strictly linked to biotransformation. For example, fluazinam and pretomanid induced significant shifts in microbial composition, characterized by the depletion of several *Clostridia* populations and the expansion of commensals, including *Akkermansia, Veillonella* and *Bifidobacterium* (Figure 2). In contrast, flutamide led to minimal detectable changes in community composition, possibly explained by its rapid biotransformation and consequently limited exposure of the parent compound to the microbial community (Figure 6). Microbial nitroreduction does not provide a nutritional benefit to the microorganisms involved.^44^ Instead, nitroreduction of drugs such as niclosamide and entacapone may serve as a “community protection” mechanism against these antibacterial compounds.^42^ Other well-documented examples include the deactivation of chloramphenicol^45^ and increased toxicity of nitrobenzodiazepines.^46^ Nitroreduction may therefore detoxify antibacterial compounds containing nitro groups, though this mechanism may not apply to flutamide, which lacks potent antibacterial properties.^47^

The identification of microbial metabolites in complex fecal matrices remains a significant challenge. To address this, we developed a computational workflow that combines biotransformation predictions (MicrobeRX and Pickaxe) with untargeted LC–MS/MS data to propose candidate microbial metabolites arising from flutamide, fluazinam, and pretomanid. By initially generating hundreds of theoretical structures and systematically filtering them through statistical analysis and visual inspection, we narrowed the field to a high-confidence subset of candidate metabolites. Following the Schymanski confidence scale, most candidate metabolites reported here correspond to level 5 (exact mass of interest), as structural annotation relied only on exact mass matching of predicted structures against MS^1^ features, without isotope pattern confirmation or MS^2^ fragmentation data.^48^ While this reflects an inherent limitation of the workflow, the proposed candidates are supported by additional evidence beyond the mass match: all structures were predicted by established biotransformation tools and were detected only in active microbiome incubations when exposed to the parent chemical and in parallel to patent chemical depletion. Although this supporting evidence is not enough to formally raise the Schymanski confidence level, it makes false annotations unlikely. The one structure formally identified with a high confidence level was flu-6, a nitroreduced metabolite of flutamide, which was purchased as a reference standard and confirmed with level 1 confidence through retention time and MS^2^ spectral matching. This metabolite was previously known to be formed in the liver^49^ but never before reported as a product of the human gut microbiome.

Nitroreduction pathways dominated the biotransformation of all three compounds, consistent with the established role of the gut microbiome as a highly reductive environment. Beyond a simple chemical transformation, nitroreduction can act as a chemical switch that alters the biological activity of the parent compounds, with potential consequences for their efficacy or toxicity. For pretomanid, the detection of the amino product (Figure 5, compound **5**) and its denitrosated derivatives (compounds **6** and **7**) confirm the capacity of the microbiota to perform the bioactivation required for therapeutic efficacy. For flutamide, among the six strains tested, we identified four strains capable of nitroreducing flutamide to the less hepatotoxic derivative flu-6. Together, these findings demonstrate how the gut microbiota can mirror or complement reductive transformations of fluorinated compounds, thereby influencing the systemic distribution of biotransformation products, with potential implications for the toxicological profiles of both clinical and environmental chemicals.

For fluazinam, depletion in abiotic controls indicates chemical instability, yet the faster degradation in microbiome incubations suggests an additional microbial contribution (Figure 4). Based on our untargeted LC-MS/MS computational pipeline analysis and prior reports, nitroreduction^50^ and hydrolytic dechlorination^51^ represent plausible biotransformation pathways, with QSAR-based predictions^51^ indicating that certain amine products may be more toxic than the parent compound, whereas hydroxylated products may be less hazardous. Pretomanid also underwent microbiome-associated nitroreduction as the key bioactivation step underlying pretomanid’s antitubercular efficacy.^52^ While reference standards would further support metabolite annotation, their limited availability precludes systematic validation.

Given that flu-6 has been reported to undergo subsequent bioactivation in other systems,^53^ its toxicological relevance warrants further consideration. Since 2-hydroxyflutamide is the pharmacologically active metabolite, further conjugation and biliary excretion raise the possibility of enterohepatic cycling and re-entry into the gut lumen, where microbial nitroreduction could further alter the overall metabolite profile. However, under the conditions tested here, flu-6 did not induce cytotoxicity in HepG2 cells even at high concentrations. These findings should be interpreted as assay- and context-dependent and do not exclude effects on other endpoints or host–microbe co-metabolic interactions that could influence therapeutic outcomes. *In silico* prediction of toxicological endpoints using the Danish QSAR database showed that flu-6 retains a similar hazard profile to flutamide, including androgen receptor antagonism, with no overall indication of additional toxicological concerns. Quantitative pharmacokinetic modelling using a microbiome-competent PBK approach further showed that systemic concentrations of both flutamide and flu-6 remained low, reaching a C_max_ of 450 nM and 33 nM, respectively, with flutamide concentrations remaining largely unchanged by microbial metabolism due to its rapid absorption via the small intestinal barrier. Although conversion of flutamide to flu-6 was complete within 1 h of fermentation, this biotransformation is unlikely to substantially affect drug efficacy or toxicity, as rapid absorption of the parent compound limits the quantitative impact of this biotransformation product on flutamide pharmacokinetics.

Together, these findings demonstrate that a subset of tested fluorinated chemicals can be metabolized by donor-derived gut microbiomes, with exposure driving compound-specific shifts in community composition. Strain-level evidence for flutamide nitroreduction provides a concrete example of how microbial metabolism may contribute to a metabolite otherwise attributed to host pathways, highlighting the role of the gut microbiota in xenobiotic metabolism. In the case of flu-6, the microbial metabolite confirmed in this study, combined *in silico* prediction of toxicological endpoints, pharmacokinetic modeling, *in vitro* cytotoxicity characterization and quantitative *in vitro* to *in vivo* extrapolation, suggests this transformation is unlikely, however, to significantly alter host effects. Finally, this work establishes an integrated *ex vivo, in vitro,* and *in silico* approach as a blueprint for quantitative and predictive strategies to understand chemical-microbiome interactions.

## Data and code availability

- PBK model parameters (including references) and code are available via Zenodo^25^
- Sequencing data generated from this study are available on the European Nucleotide Archive (ENA) under the study accession PRJEB112299. Code used for corresponding analysis is available on GitHub (https://github.com/jorgepena9/fluxigut-analysis).
- LC-MS/MS data and data analysis files are available for reviewers on Metabolights and will be made public upon acceptance of the manuscript.

## Acknowledgements

The authors acknowledge funding support from ETH Research Grants (23-2 ETH-047, to N.A.B., S.J.S., and S.L.R.) and the Uniscientia Stiftung (to N.A.B. and S.L.R.). S.I.P. is supported by the Peter und Traudl Engelhorn Foundation.

## Notes

### Competing Interest Statement

The authors have declared no competing interest.

## References

1. Inoue, M., Sumii, Y., and Shibata, N. (2020). Contribution of Organofluorine Compounds to Pharmaceuticals. ACS Omega 5, 10633–10640. 10.1021/acsomega.0c00830.

2. Ogawa, Y., Tokunaga, E., Kobayashi, O., Hirai, K., and Shibata, N. (2020). Current Contributions of Organofluorine Compounds to the Agrochemical Industry. iScience 23, 101467. 10.1016/j.isci.2020.101467.

3. Johnson, B.M., Shu, Y.Z., Zhuo, X., and Meanwell, N.A. (2020). Metabolic and Pharmaceutical Aspects of Fluorinated Compounds. J Med Chem 63, 6315–6386. 10.1021/acs.jmedchem.9b01877.

4. Gluge, J., Scheringer, M., Cousins, I.T., DeWitt, J.C., Goldenman, G., Herzke, D., Lohmann, R., Ng, C.A., Trier, X., and Wang, Z. (2020). An overview of the uses of per- and polyfluoroalkyl substances (PFAS). Environ Sci Process Impacts 22, 2345–2373. 10.1039/d0em00291g.

5. Probst, S.I., Felder, F.D., Poltorak, V., Mewalal, R., Blaby, I.K., and Robinson, S.L. (2025). Enzymatic carbon-fluorine bond cleavage by human gut microbes. Proc Natl Acad Sci U S A 122, e2504122122. 10.1073/pnas.2504122122.

6. Koppel, N., Rekdal, V.M., and Balskus, E.P. (2017). Chemical transformation of xenobiotics by the human gut microbiota. Science 356, eaag2770. 10.1126/science.aag2770.

7. Lehouritis, P., Cummins, J., Stanton, M., Murphy, C.T., McCarthy, F.O., Reid, G., Urbaniak, C., Byrne, W.L., and Tangney, M. (2015). Local bacteria affect the efficacy of chemotherapeutic drugs. Sci Rep 5, 14554. 10.1038/srep14554.

8. Spanogiannopoulos, P., Kyaw, T.S., Guthrie, B.G.H., Bradley, P.H., Lee, J.V., Melamed, J., Malig, Y.N.A., Lam, K.N., Gempis, D., Sandy, M., et al. (2022). Host and gut bacteria share metabolic pathways for anti-cancer drug metabolism. Nat Microbiol 7, 1605–1620. 10.1038/s41564-022-01226-5.

9. R Okeda, M.S., T Matsuo, T Kuroiwa, R Shimokawa, T Tajima (1990). Experimental neurotoxicity of 5-fluorouracil and its derivatives is due to poisoning by the monofluorinated organic metabolites, monofluoroacetic acid and alpha-fluoro-beta-alanine. Acta Neuropathol 81, 66–73. 10.1007/BF00662639.

10. Franck, C., Malfertheiner, P., and Venerito, M. (2017). Safe administration of S-1 after 5-fluorouracil-induced cardiotoxicity in a patient with colorectal cancer. BMJ Case Rep 2017. 10.1136/bcr-2016-219162.

11. Locke MA, Z.R., Steinriede RW, Kingery WL. (2007). Degradation and sorption of fluometuron and metabolites in conservation tillage soils. Journal of Agricultural and Food Chemistry 55, 844–851.

12. Gopal, A., Swamidason, J., Mariappan, P., and Bojan, V. (2025). Microbial degradation of flubendiamide in different types of soils at tropical region using lactic acid bacteria formulation. Sci Rep 15, 29271. 10.1038/s41598-025-08917-z.

13. Mastrorilli, E., Herd, P., Rey, F.E., Goodman, A.L., and Zimmermann, M. (2026). Linking interpersonal differences in gut microbiota composition and drug biotransformation activity. bioRxiv. 10.64898/2026.01.21.700809.

14. Zund, J.N., Pluss, S., Mujezinovic, D., Menzi, C., von Bieberstein, P.R., de Wouters, T., Lacroix, C., Leventhal, G.E., and Pugin, B. (2024). A flexible high-throughput cultivation protocol to assess the response of individuals’ gut microbiota to diet-, drug-, and host-related factors. ISME Commun 4, ycae035. 10.1093/ismeco/ycae035.

15. Mendez-Catala, D.M., Spenkelink, A., Rietjens, I., and Beekmann, K. (2020). An in vitromodel to quantify interspecies differences in kinetics for intestinal microbial bioactivation and detoxification of zearalenone. Toxicol Rep 7, 938–946. 10.1016/j.toxrep.2020.07.010.

16. Ruiz-Moreno, A.J., Del Castillo-Izquierdo, A., Tamargo-Rubio, I., and Fu, J. (2025). MicrobeRX: a tool for enzymatic-reaction-based metabolite prediction in the gut microbiome. Microbiome 13, 78. 10.1186/s40168-025-02070-5.

17. Shebek, K.M., Strutz, J., Broadbelt, L.J., and Tyo, K.E.J. (2023). Pickaxe: a Python library for the prediction of novel metabolic reactions. BMC Bioinformatics 24, 106. 10.1186/s12859-023-05149-8.

18. Stevanoska, M., Cremona, M., Beekmann, K., Sturla, S.J., and Aichinger, G. (2026). Interindividual variability in gut microbial formation of the hop phytoestrogen 8-prenylnaringenin results in elevated but sub-toxic internal exposures. Arch Toxicol. 10.1007/s00204-026-04329-8.

19. Punt, A., Pinckaers, N., Peijnenburg, A., and Louisse, J. (2021). Development of a web-based toolbox to support quantitative in-vitro-to-in-vivo extrapolations (QIVIVE) within nonanimal testing strategies. Chem Res Toxicol 34, 460–472. 10.1021/acs.chemrestox.0c00307.

20. Cheng T, Z.Y., Li X, Lin F, Xu Y, Zhang X, Li Y, Wang R, Lai L. (2007). Computation of octanol−water partition coefficients by guiding an additive model with knowledge. J Chem Inf Model 26, 2140–2148.

21. van Tongeren, T.C.A., Carmichael, P.L., Rietjens, I., and Li, H. (2022). Next Generation Risk Assessment of the Anti-Androgen Flutamide Including the Contribution of Its Active Metabolite Hydroxyflutamide. Front Toxicol 4, 881235. 10.3389/ftox.2022.881235.

22. Kamiya, Y., Takaku, H., Yamada, R., Akase, C., Abe, Y., Sekiguchi, Y., Murayama, N., Shimizu, M., Kitajima, M., Shono, F., et al. (2020). Determination and prediction of permeability across intestinal epithelial cell monolayer of a diverse range of industrial chemicals/drugs for estimation of oral absorption as a putative marker of hepatotoxicity. Toxicol Rep 7, 149– 154. 10.1016/j.toxrep.2020.01.004.

23. Sun D, L.H., Welage L S, Barnett J L, Landowski C P, Foster D, Fleisher D, Lee K D, and Amidon G L (2002). Comparison of human duodenum and Caco-2 gene expression profiles for 12,000 gene sequences tags and correlation with permeability of 26 drugs. Pharmacol Res 19, 1400–1416.

24. Radwanski, E., Perentesis, G., Symchowicz, S., and Zampaglione, N. (1989). Single and multiple dose pharmacokinetic evaluation of flutamide in normal geriatric volunteers. J Clin Pharmacol 29, 554–558. 10.1002/j.1552-4604.1989.tb03381.x.

25. Aichinger (2026). Zenodo. 10.5281/zenodo.19682173.

26. Yu, Y., Trottmann, N.F., Scharer, M.R., Fenner, K., and Robinson, S.L. (2024). Substrate promiscuity of xenobiotic-transforming hydrolases from stream biofilms impacted by treated wastewater. Water Res 256, 121593. 10.1016/j.watres.2024.121593.

27. Heuckeroth, S., Damiani, T., Smirnov, A., Mokshyna, O., Brungs, C., Korf, A., Smith, J.D., Stincone, P., Dreolin, N., Nothias, L.F., et al. (2024). Reproducible mass spectrometry data processing and compound annotation in MZmine 3. Nat Protoc 19, 2597–2641. 10.1038/s41596-024-00996-y.

28. Flörl L, C.P., Moccia MD, Plüss S, Bokulich NA. (2026). HighALPS ultra-high-throughput marker-gene amplicon library preparation and sequencing on the Illumina NextSeq and NovaSeq Platforms. Msystems, 00023–00026.

29. Apprill, A., McNally, S., Parsons, R., and Weber, L. (2015). Minor revision to V4 region SSU rRNA 806R gene primer greatly increases detection of SAR11 bacterioplankton. Aquat Microb Ecol 75, 129–137. 10.3354/ame01753.

30. Parada, A.E., Needham, D.M., and Fuhrman, J.A. (2016). Every base matters: assessing small subunit rRNA primers for marine microbiomes with mock communities, time series and global field samples. Environ Microbiol 18, 1403–1414. 10.1111/1462-2920.13023.

31. Bolyen, E., Rideout, J.R., Dillon, M.R., Bokulich, N.A., Abnet, C.C., Al-Ghalith, G.A., Alexander, H., Alm, E.J., Arumugam, M., Asnicar, F., et al. (2019). Reproducible, interactive, scalable and extensible microbiome data science using QIIME 2. Nat Biotechnol 37, 852–857. 10.1038/s41587-019-0209-9.

32. Callahan, B.J., McMurdie, P.J., Rosen, M.J., Han, A.W., Johnson, A.J., and Holmes, S.P. (2016). DADA2: High-resolution sample inference from Illumina amplicon data. Nat Methods 13, 581–583. 10.1038/nmeth.3869.

33. Kaehler, B.D., Bokulich, N.A., McDonald, D., Knight, R., Caporaso, J.G., and Huttley, G.A. (2019). Species abundance information improves sequence taxonomy classification accuracy. Nat Commun 10, 4643. 10.1038/s41467-019-12669-6.

34. Bokulich, N.A., Kaehler, B.D., Rideout, J.R., Dillon, M., Bolyen, E., Knight, R., Huttley, G.A., and Gregory Caporaso, J. (2018). Optimizing taxonomic classification of marker-gene amplicon sequences with QIIME 2’s q2-feature-classifier plugin. Microbiome 6, 90. 10.1186/s40168-018-0470-z.

35. Robeson, M.S., 2nd, O’Rourke, D.R., Kaehler, B.D., Ziemski, M., Dillon, M.R., Foster, J.T., and Bokulich, N.A. (2021). RESCRIPt: Reproducible sequence taxonomy reference database management. PLoS Comput Biol 17, e1009581. 10.1371/journal.pcbi.1009581.

36. Chuvochina, M., Gerken, J., Frentrup, M., Sandikci, Y., Goldmann, R., Freese, H.M., Goker, M., Sikorski, J., Yarza, P., Quast, C., et al. (2026). SILVA in 2026: a global core biodata resource for rRNA within the DSMZ digital diversity. Nucleic Acids Res 54, D334–D341. 10.1093/nar/gkaf1247.

37. Lin, H., and Peddada, S.D. (2024). Multigroup analysis of compositions of microbiomes with covariate adjustments and repeated measures. Nat Methods 21, 83–91. 10.1038/s41592-023-02092-7.

38. Roux, I., Lindell, A.E., Griesshammer, A., Smith, T., Krishna, S., Guan, R., Rad, D., Faria, L., Blasche, S., Patil, K.R., et al. (2025). Industrial and agricultural chemicals exhibit antimicrobial activity against human gut bacteria in vitro. Nat Microbiol 10, 3107–3121. 10.1038/s41564-025-02182-6.

39. Zimmermann, M., Zimmermann-Kogadeeva, M., Wegmann, R., and Goodman, A.L. (2019). Mapping human microbiome drug metabolism by gut bacteria and their genes. Nature 570, 462–467. 10.1038/s41586-019-1291-3.

40. Wedebye, E.B., Dybdahl, M., Reffstrup, T.K., Rosenberg, S.A., Løfstedt, M., and Nikolov, N.G. (2016). The new Danish (Q)SAR database: A freely available tool with predictions for >600,000 substances. Toxicol Lett 258, S118. 10.1016/j.toxlet.2016.06.1479.

41. Lindell, A.E., Griesshammer, A., Michaelis, L., Papagiannidis, D., Ochner, H., Kamrad, S., Guan, R., Blasche, S., Ventimiglia, L.N., Ramachandran, B., et al. (2025). Human gut bacteria bioaccumulate per- and polyfluoroalkyl substances. Nat Microbiol 10, 1630–1647. 10.1038/s41564-025-02032-5.

42. Garcia-Santamarina, S., Kuhn, M., Devendran, S., Maier, L., Driessen, M., Mateus, A., Mastrorilli, E., Brochado, A.R., Savitski, M.M., Patil, K.R., et al. (2024). Emergence of community behaviors in the gut microbiota upon drug treatment. Cell 187, 6346–6357 e6320. 10.1016/j.cell.2024.08.037.

43. Penning, T.M., Su, A.L., and El-Bayoumy, K. (2022). Nitroreduction: A Critical Metabolic Pathway for Drugs, Environmental Pollutants, and Explosives. Chem Res Toxicol 35, 1747–1765. 10.1021/acs.chemrestox.2c00175.

44. Peres CM, A.S., Agathos, S.N. (2000). Biodegradation of nitroaromatic pollutants from pathways to remediation. Biotechnol Annu Rev 6, 197–200. 10.1016/s1387-2656(00)06023-3

45. Crofts, T.S., Sontha, P., King, A.O., Wang, B., Biddy, B.A., Zanolli, N., Gaumnitz, J., and Dantas, G. (2019). Discovery and Characterization of a Nitroreductase Capable of Conferring Bacterial Resistance to Chloramphenicol. Cell Chem Biol 26, 559–570.e556. 10.1016/j.chembiol.2019.01.007.

46. Linwu, S.W., Syu, C.J., Chen, Y.L., Wang, A.H., and Peng, F.C. (2009). Characterization of Escherichia coli nitroreductase NfsB in the metabolism of nitrobenzodiazepines. Biochem Pharmacol 78, 96–103. 10.1016/j.bcp.2009.03.019.

47. Kruszewska, H., Zareba, T., and Tyski, S. (2008). Examination of antibacterial and antifungal activity of selected non-antibiotic products. Acta Pol Pharm 65, 779–782.

48. Schymanski, E.L., Jeon, J., Gulde, R., Fenner, K., Ruff, M., Singer, H.P., and Hollender, J. (2014). Identifying small molecules via high resolution mass spectrometry: communicating confidence. Environ Sci Technol 48, 2097–2098. 10.1021/es5002105.

49. Kang, P., Dalvie, D., Smith, E., Zhou, S., Deese, A., and Nieman, J.A. (2008). Bioactivation of flutamide metabolites by human liver microsomes. Drug Metab Dispos 36, 1425–1437. 10.1124/dmd.108.020370.

50. Kolanczyk, R.C., Serrano, J.A., Tapper, M.A., and Schmieder, P.K. (2018). A comparison of fish pesticide metabolic pathways with those of the rat and goat. Regul Toxicol Pharmacol 94, 124–143. 10.1016/j.yrtph.2018.01.019.

51. Li, N., Xia, Y., Li, Y., Jia, Q., Qiu, J., Xu, Y., Wang, Z., Liu, Z., and Qian, Y. (2023). Untargeted screening, quantitative analysis, and toxicity estimation of degradation products of fluazinam in vegetables. Microchem J 190, 108584. 10.1016/j.microc.2023.108584.

52. EMA (2020). EMA Assessment report Pretomanid 2020. Contract No.: EMEA/H/C/005167/0000.

53. Wen B, C.K., Rademacher P, Fitch WL, Monshouwer M, Nelson SD. (2008). Comparison of in Vitro Bioactivation of Flutamide and Its Cyano Analogue Evidence for Reductive Activation by Human NADPH Cytochrome P450 Reductase. Chem Res Toxicol 15, 2393–2406. 10.1021/tx800281h.

