## Supplementary Information for "Bidirectional interactions between gut microbiota and fluorochemical biotransformation and bioactivity"

**Table S1**. Basal yeast casitone fatty acid (bYCFA), short chain fatty acid (SCFA) and complex carbohydrate and mucin (6C+Muc) media components.

|  | **Chemical** | **Vendor** |
| --- | --- | --- |
|  | Amicase | Sigma Aldrich |
|  | Yeast extract | Sigma Aldrich |
|  | Meat extract | Sigma Aldrich |
|  | K_2_HPO_4_ | VWR Life Science |
|  | KH_2_PO_4_ | VWR Life Science |
|  | NaCl | Sigma Aldrich |
| bYCFA | (NH_4_)_2_SO_4_ | Sigma Aldrich |
|  | MgSO_4_ (*7H_2_O) | Sigma Aldrich |
|  | CaCl2 (*2H_2_O) | VWR Life Science |
|  | _L_-cysteine HCl (*H_2_O) | VWR Chemicals |
|  | NaHCO_3_ | Merck |
|  | Hemin | Across Chemicals |
|  | Resazurin | Sigma Aldrich |
|  | Acetic acid | Merck |
|  | Propionic acid | Sigma Aldrich |
| SCFA | Valeric acid | Thermo Scientific |
|  | Isovaleric acid | ACROS Organics |
|  | Isobutyric acid | ACROS Organics |
|  | Starch | Sigma Aldrich |
|  | Pectin | Alpha Aesar |
|  | Xylan | Apollo Scientific |
| 6CMuc | Arabinogalactan | Sigma Aldrich |
|  | Guar | Sigma Aldrich |
|  | Inulin | Sigma Aldrich |
|  | Mucin type II | Sigma Aldrich |

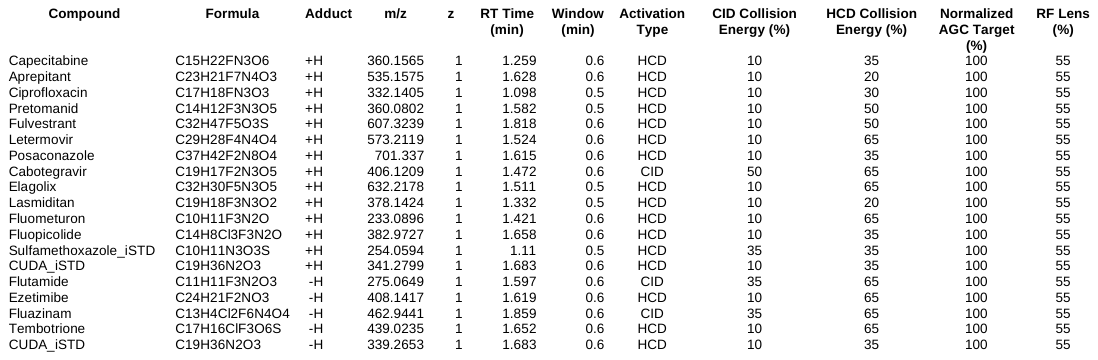
**Table S2.** Transition list used for targeted metabolomics in Skyline.

**Table S3**. Untargeted metabolomic data processing parameters in MSDIAL version 4.9.

| MS1 Data type | Centroid | | Centroid |
| --- | --- | --- | --- |
| MS2 Data type | Centroid | | Centroid |
| Ion mode | Positive | | Negative |
| Target | Metablomics | | Metablomics |
| Mode | ddMSMS | | ddMSMS |
| #Data collection parameters |  | |  |
| Retention time begin | 0 | | 0 |
| Retention time end | 100 | | 100 |
| Mass range begin | 0 | | 0 |
| Mass range end | 2000 | | 2000 |
| MS2 mass range begin | 0 | | 0 |
| MS2 mass range end | 2000 | | 2000 |
| #Centroid parameters |  | |  |
| MS1 tolerance | 0.008 | | 0.008 |
| MS2 tolerance | 0.025 | | 0.025 |
| #Isotope recognition |  | |  |
| Maximum charged number | 2 | | 2 |
| #Data processing |  | |  |
| Number of threads | 11 | | 1 |
| #Peak detection parameters |  | |  |
| Smoothing method | LinearWeightedMovingAverage | | LinearWeightedMovingAverage |
| Smoothing level | 3 | | 3 |
| Minimum peak width | 6 | | 5 |
| Minimum peak height | 80000 | | 40000 |
| #Peak spotting parameters |  | |  |
| Mass slice width | 0.1 | | 0.1 |
| Exclusion mass list (mass & tolerance) |  | |  |
| #Deconvolution parameters |  | |  |
| Sigma window value | 0.5 | | 0.5 |
| MS2Dec amplitude cut off | 0 | | 0 |
| Exclude after precursor | TRUE | | TRUE |
| Keep isotope until | 0.5 | | 0.5 |
| Keep original precursor isotopes | FALSE | | FALSE |
| #MSP file and MS/MS identification setting |  | |  |
| MSP file | MoNA_NIST23_wInSilico_POS.msp | | MoNA_NIST23_wInSilico_NEG.msp |
| Retention time tolerance | 100 | | 100 |
| Accurate mass tolerance (MS1) | 0.01 | | 0.01 |
| Accurate mass tolerance (MS2) | 0.1 | | 0.1 |
| Identification score cut off | 80 | | 80 |
| Using retention time for scoring | FALSE | | FALSE |
| Using retention time for filtering | FALSE | | FALSE |
| #Post identification (retention time and accurate mass based) setting |  | |  |
| Retention time tolerance | 0.15 | | 0.15 |
| Identification score cut off | 85 | | 85 |
| #Advanced setting for identification |  | |  |
| Relative abundance cut off | 0 | | 0 |
| Top candidate report | FALSE | | FALSE |
| #Adduct ion setting | [M+H]+ | | [M-H]- |
|  | [M+NH4]+ | | [M+Cl]- |
|  | [M+Na]+ | | [2M-H]- |
|  | [2M+H]+ | |  |
|  | [2M+NH4]+ | |  |
|  | [2M+Na]+ | |  |
| #Alignment parameters setting |  | |  |
| Reference file | 250815_QBS_Blank_26_RPpos.raw | | 250815_QBS_QC_26_RPneg.raw |
| Retention time tolerance | 0.1 | | 0.07 |
| MS1 tolerance | 0.015 | | 0.015 |
| Retention time factor | 0.5 | | 0.5 |
| MS1 factor | 0.5 | | 0.5 |
| Peak count filter | 0 | | 0 |
| N% detected in at least one group | 0 | | 0 |
| Remove feature based on peak height fold-change | TRUE | | TRUE |
| Sample max / blank average | 5 | | 5 |
| Sample average / blank average | 5 | | 5 |
| Keep identified and annotated metabolites | TRUE | | TRUE |
| Keep removable features and assign the tag for checking | TRUE | | TRUE |
| Gap filling by compulsion | TRUE | | TRUE |
| #Tracking of isotope labels |  | |  |
| Tracking of isotopic labels | FALSE | | FALSE |
| #Ion mobility |  | |  |
| Ion mobility data | FALSE | | FALSE |

**Table S4.** Proposed microbially-derived metabolites of flutamide, fluazinam, and pretomanid.

| Parent | Compound | Formula | Ret. time (min) | Measured *m/z* | Predicted *m/z* | Theoretical adduct | Fold change | MWU  p-value |
| --- | --- | --- | --- | --- | --- | --- | --- | --- |
| Flutamide | **1** | C_11_H_11_F_3_N_2_O_2_ | 1.236 | 259.06976 | 259.069986 | [M-H]^-^ | 0.466 | 9.06E-06 |
|  | Flu-6 | C_11_H_13_F_3_N_2_O | 1.290 | 247.10503 | 247.105273 | [M+H] ^+^ | 7.686 | 1.94E-53 |
| Fluazinam | **2** | C_13_H_6_Cl_2_F_6_N_3_O_2_ | 1.721 | 434.98462 | 434.984476 | [M+H] ^+^ | 9.316 | 2.24E-203 |
|  | **3** | C_13_H_8_Cl_2_F_6_N_3_ | 1.664 | 405.01068 | Not predicted | [M+H] ^+^ | inf | 1.23E-45 |
|  | **4** | C_13_H_5_ClF_6_N_4_O_5_ | 1.622 | 444.97784 | 444.977990 | [M-H] ^-^ | inf | 1.25E-116 |
| Pretomanid | **5** | C_14_H_14_F_3_N_3_O_3_ | 1.049 | 347.13138 | 347.13217 | [M+NH_4_] ^+^ | 4.6148 | 6.23E-70 |
|  | **6** | C_14_H_13_F_3_N_2_O_4_ | 1.040 | 348.13318 | 348.116193 | [M+NH_4_] ^+^ | 9.085 | 4.55E-175 |
|  | **7** | C_14_H_13_F_3_N_2_O_5_ | 1.082 | 364.11185 | 364.111108 | [M+NH_4_] ^+^ | 9.700 | 2.12E-221 |
|  | **8** | C_15_H_15_N_3_O_6_ | 1.294 | 332.08594 | 332.088811 | [M-H] ^-^ | 9.208 | 4.16E-225 |

**Table S5**. Reduction of the potential gut bacterial metabolic candidates of flutamide, fluazinam, and pretomanid after every step of the computational identification pipeline.

|  | **Flutamide** | **Fluazinam** | **Pretomanid** |
| --- | --- | --- | --- |
| Initially predicted metabolic structures | 471 | 268 | 431 |
| Unique molecular formulas of predictions | 152 | 83 | 120 |
| After matching *m/z* untargeted values | 742 | 225 | 958 |
| After statistics-based filtering | 41 | 24 | 49 |
| After visual inspection of formation trend | 2 | 2 | 4 |

***Table S6****. QSAR predictions for flutamide and flu-6*

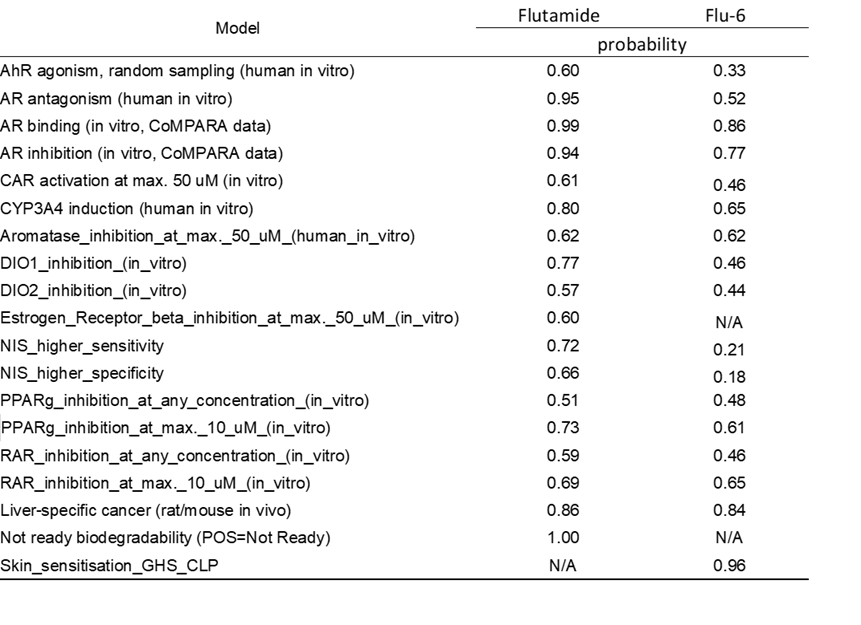

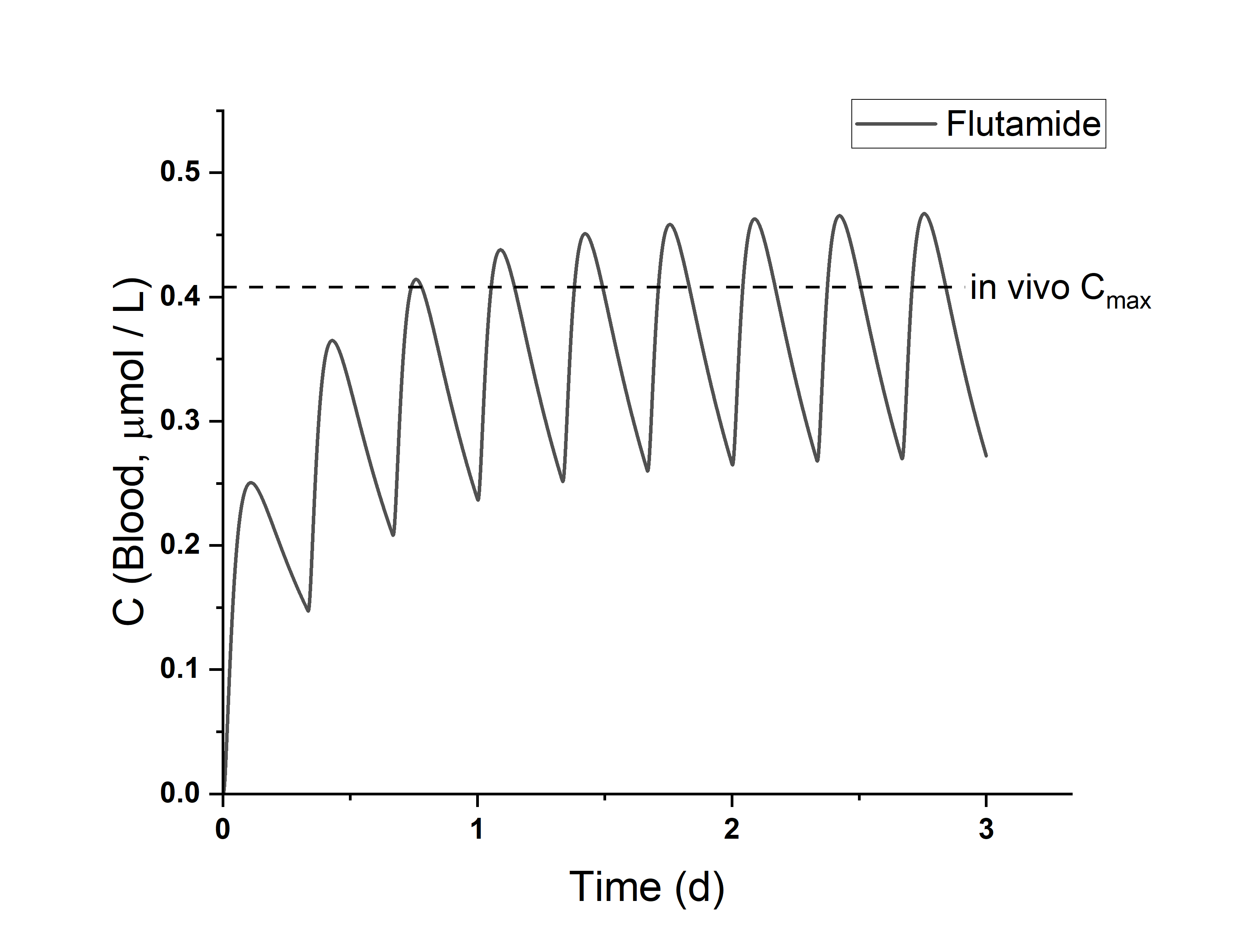

**Figure S1. PBK model evaluation.** PBK-model predicted blood concentrations of flutamide over time (gray line), assuming an intake of 250 mg of flutamide, 3 times per day, in comparison to clinical pharmacokinetics (blood steady-state C_max_ levels) from Radwanski et al. (J Clin Pharmacol, 1989).^24^

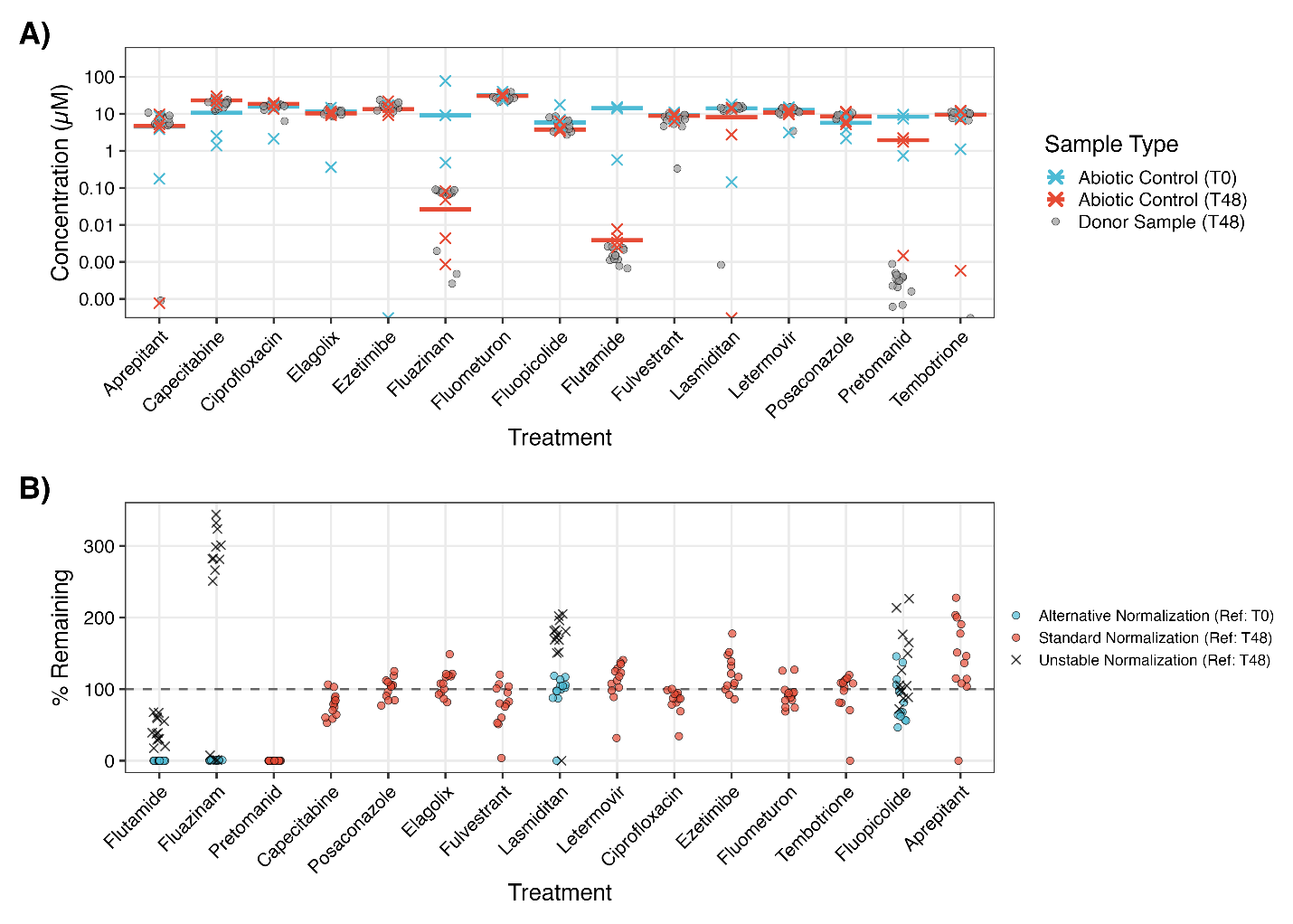

**Figure S2. Fluorinated compound stability and normalization. A)** Comparison of compound concentrations in vehicle controls (T0 and T48) and donor samples collected 48 h post-inoculation. Bars represent median values. **B)** Percent of compound remaining in donor samples at 48 h post-inoculation. Stable compounds (red circles) were normalized to the matched T48 abiotic control. Compounds exhibiting vehicle instability (blue circles) were normalized to the baseline (T0) concentrations. Black “x” markers indicate uncorrected values for vehicle-unstable compounds.

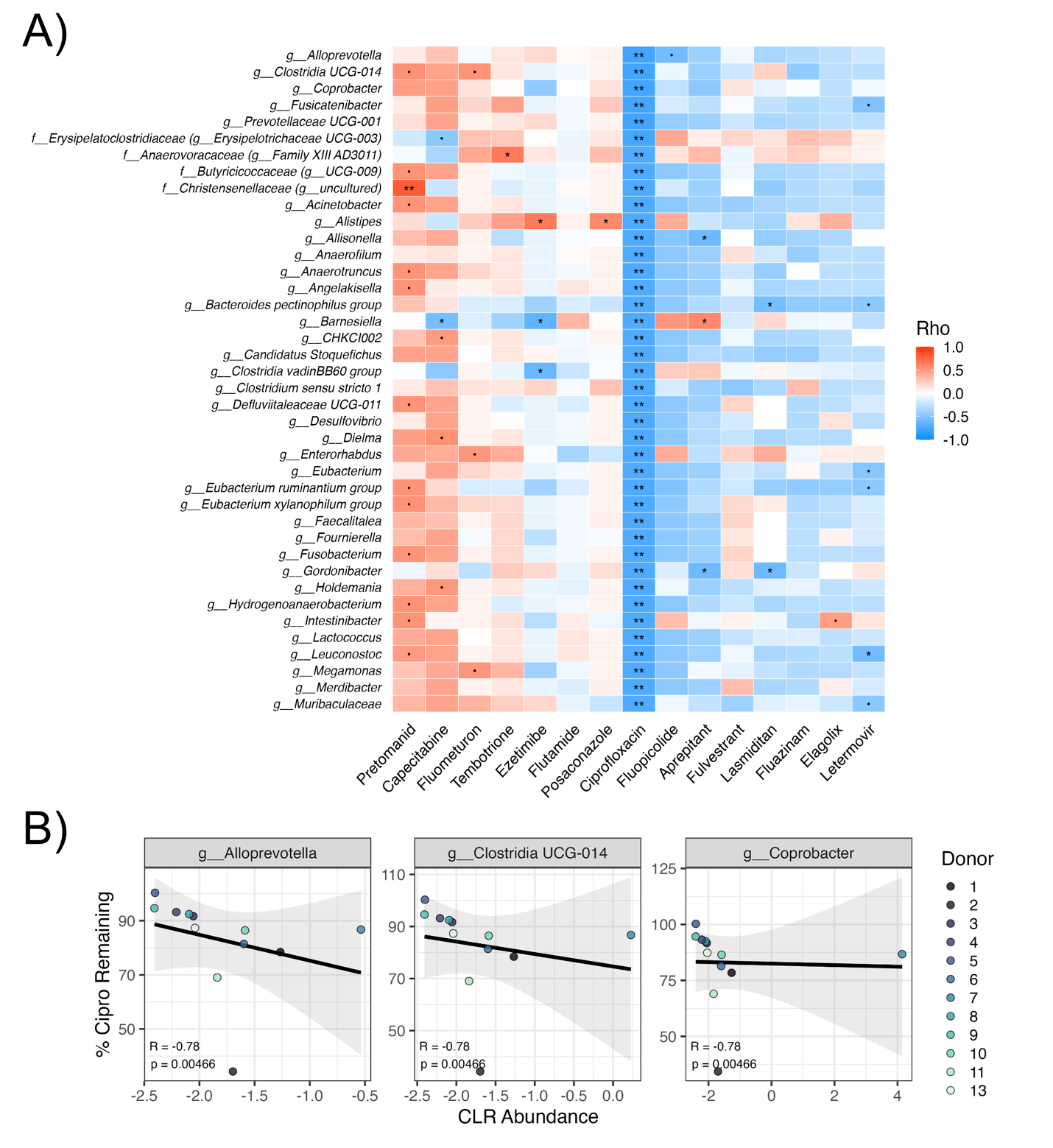

**Figure S3. Ciprofloxacin drives negative correlations through broad-spectrum antimicrobial activity.**

**A)** Heatmap of Spearman rank correlation coefficients between centered log-ratio (CLR) transformed bacterial abundances and the normalized percentage of drug remaining at 48 h post-inoculation (Figure S1). The heatmap displays the top 40 genera most negatively correlated with ciprofloxacin. The distinct, strong negative correlation pattern observed with ciprofloxacin highlights the broad-spectrum antimicrobial susceptibility of the treatment. **B)** Scatter plots of three representative taxa. Negative correlations likely reflect antimicrobial activity rather than biotransformation. Points represent individual samples colored by donor. Black lines indicate a linear regression with 95% confidence intervals.

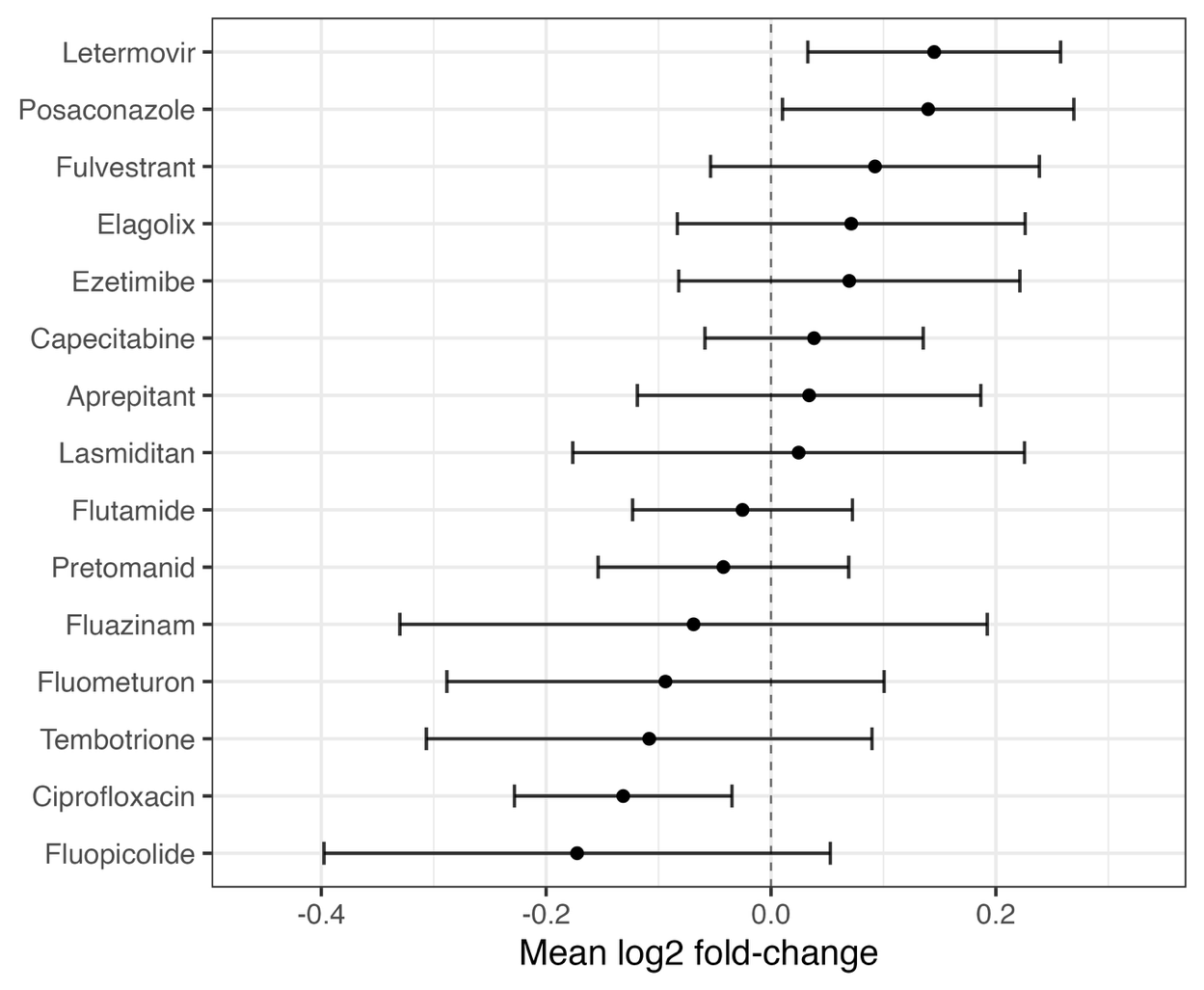

**Figure S4. Optical density remains stable across treatments.**

Mean log_2_ fold-change in bacterial optical density (OD_600_) at 48 h relative to the matched vehicle controls. Points represent the mean change across all donors, and error bars indicate 95% confidence intervals. No compounds were significant after Benjamini–Hochberg false discovery rate correction (q > 0.05).

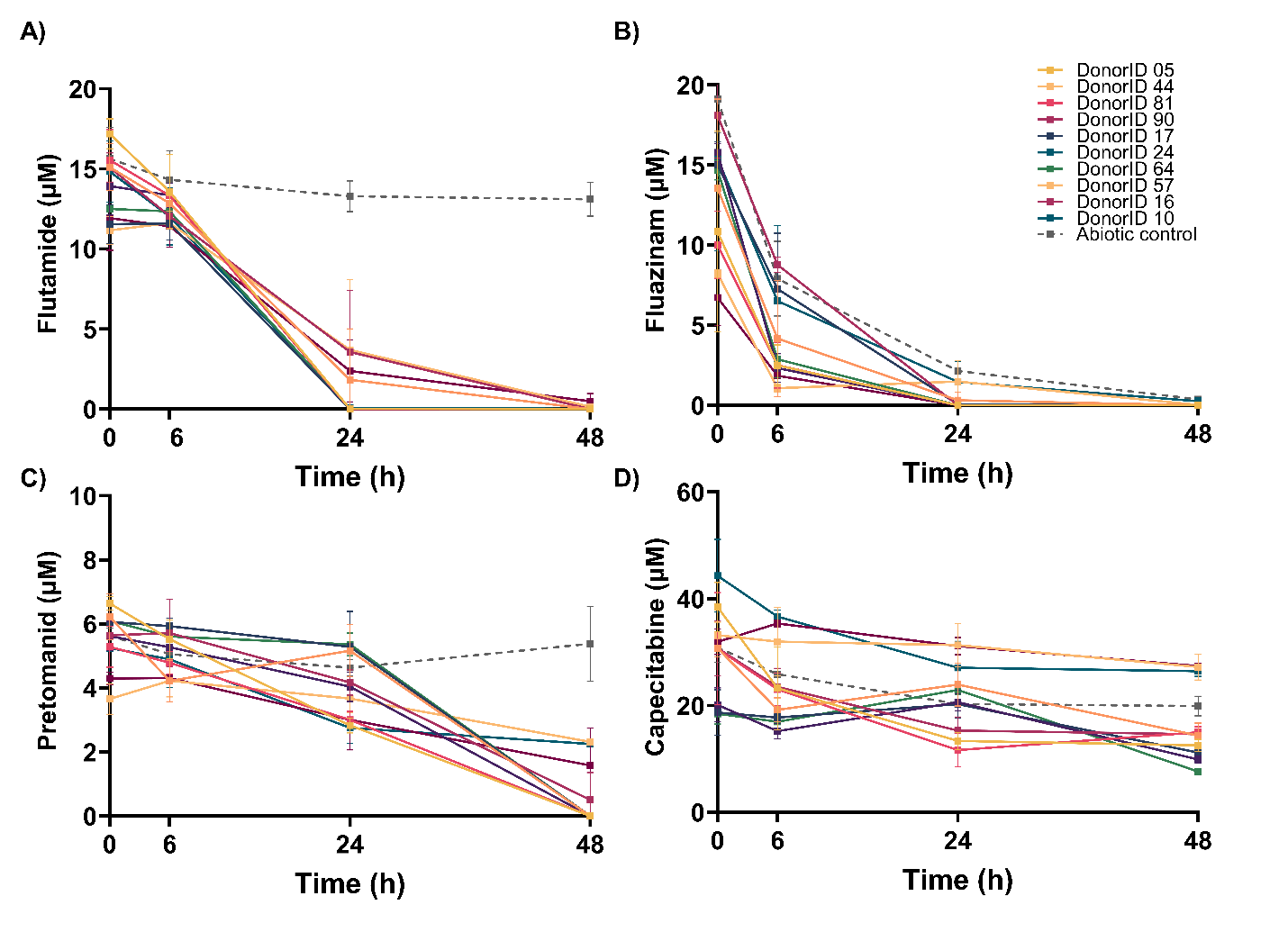

**Figure S5**. Kinetics of microbial metabolism of **A)** flutamide, **B)** fluazinam, **C)** pretomanid, and **D)** capecitabine over time across 10 donors. Dashed lines represent the abiotic controls. Data are represented as mean ± SD of three technical replicates.

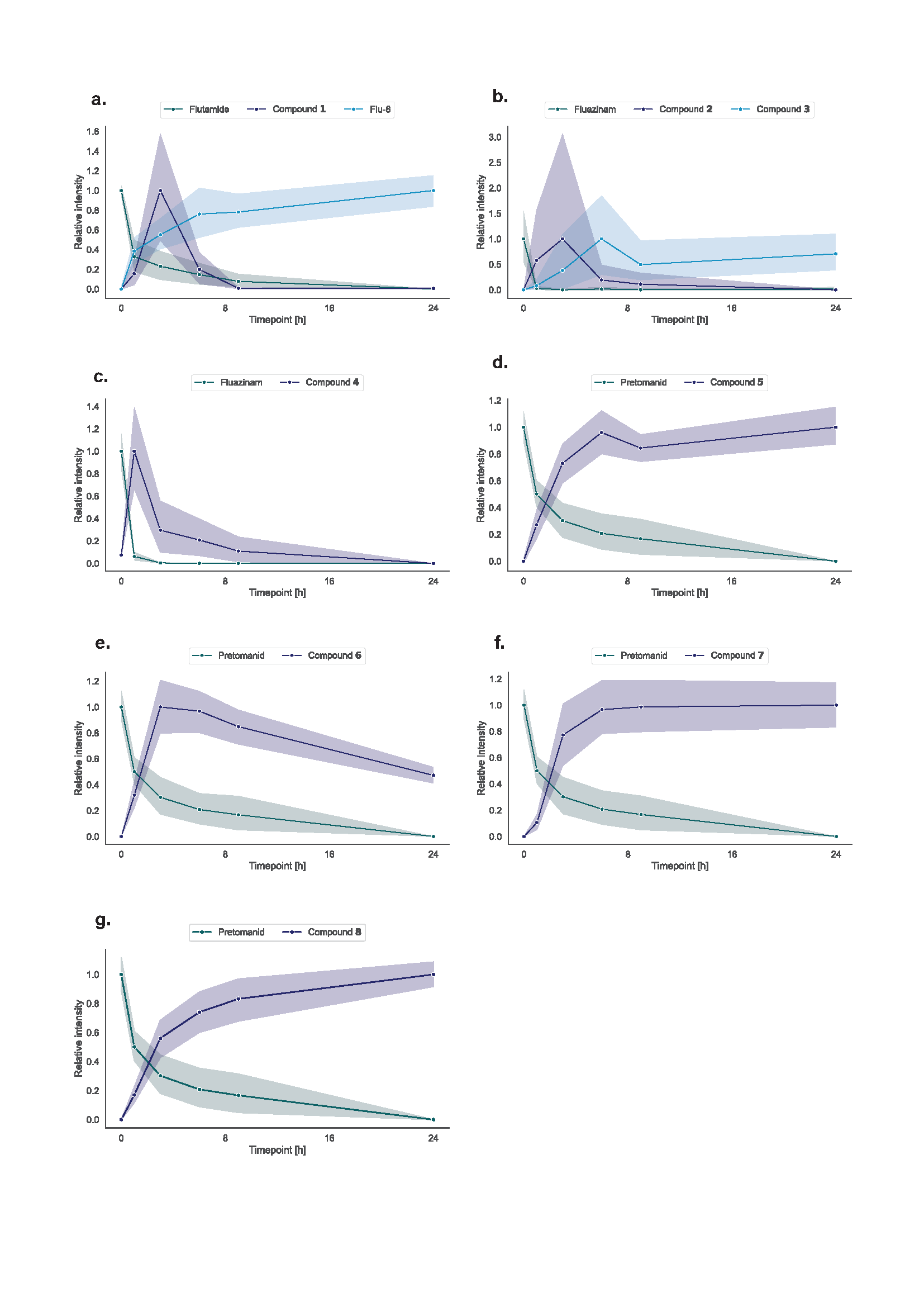

**Figure S6.** Formation trend over time for (a) compound **1** and flu-6, (b) compounds **2** and **3**, (c) compound **4**, (d) compound **5**, (e) compound **6**, (f) compound **7**, and (g) compound **8***.*
